# Comparative analysis of machine learning and evolutionary optimization algorithms for precision tissue culture of *Cannabis sativa*: Prediction and validation of *in vitro* shoot growth and development based on the optimization of light and carbohydrate sources

**DOI:** 10.1101/2021.08.09.455719

**Authors:** Marco Pepe, Mohsen Hesami, Finlay Small, Andrew Maxwell Phineas Jones

**Affiliations:** Gosling Research Institute for Plant Preservation, Department of Plant Agriculture, University of Guelph, Guelph, ON, N1G 2W1, Canada; Department of Research and Development, Entourage Health Corp, Guelph, Ontario, Canada

**Keywords:** artificial neural networks, biogeography-based optimization, genetic algorithm, interior search algorithm, *in vitro* culture, neuro-fuzzy logic, symbiotic organisms search

## Abstract

Micropropagation techniques offer opportunity to proliferate, maintain, and study dynamic plant responses in highly controlled environments without confounding external influences, forming the basis for many biotechnological applications. With medicinal and recreational interests for *Cannabis sativa* L. growing, research related to the optimization of *in vitro* practices is needed to improve current methods while boosting our understanding of the underlying physiological processes. Unfortunately, due to the exorbitantly large array of factors influencing tissue culture, existing approaches to optimize *in vitro* methods are tedious and time-consuming. Therefore, there is great potential to use new computational methodologies for analysing data to develop improved protocols more efficiently. Here, we first tested the effects of light qualities using assorted combinations of Red, Blue, Far Red, and White spanning 0-100 μmol/m^2^/s in combination with sucrose concentrations ranging from 1-6 % (w/v), totaling 66 treatments, on *in vitro* shoot growth, root development, number of nodes, shoot emergence, and canopy surface area. Collected data were then assessed using multilayer perceptron (MLP), generalized regression neural network (GRNN), and adaptive neuro-fuzzy inference system (ANFIS) to model and predict *in vitro Cannabis* growth and development. Based on the results, GRNN had better performance than MLP or ANFIS and was consequently selected to link different optimization algorithms (genetic algorithm, biogeography-based optimization, interior search algorithm, and symbiotic organisms search) for prediction of optimal light levels (quality/intensity) and sucrose concentration for various applications. Predictions of *in vitro* conditions to refine growth responses were subsequently tested in a validation experiment and data showed no significant differences between predicted optimized values and observed data. Thus, this study demonstrates the potential of machine learning and optimization algorithms to predict the most favourable light combinations and sucrose levels to elicit specific developmental responses. Based on these, recommendations of light and carbohydrate levels to promote specific developmental outcomes for *in vitro Cannabis* are suggested. Ultimately, this work showcases the importance of light quality and carbohydrate supply in directing plant development as well as the power of machine learning approaches to investigate complex interactions in plant tissue culture.

## Introduction

The multifaceted value of *Cannabis sativa* L. (cannabis) as a quality fiber, seed oil, and therapeutic crop have been recognized for millennia (Hesami et al., 2020; Sandler et al., 2019). Over the past two decades, interest relating to its medicinal applications have largely been emphasized due to the discovery of over 500 unique secondary metabolites (ElSohly and Gul, 2014). Of these compounds, there are more than 100 cannabinoids that contribute to cannabis’ pharmacological properties (Fathordoobady et al., 2019). Medicinal use can relieve symptoms associated with glaucoma, nausea, irritability, epilepsy, chronic pain, etc. (Barrus et al., 2017), showing potential to revolutionize the pharmaceutical industry, and technologies related to extraction and administration of bioactive compounds (Fathordoobady et al., 2019; Vita et al., 2020). Due to the important industrial implications of drug-type cannabis, it is imperative to establish methods for the production of high quality biomass with consistent secondary metabolite profiles, which is achievable in part through micropropagation (Chandra et al., 2020).

Since many nations have adopted the more liberal view of cannabis, it’s since gained higher economic status as an industrial crop, and additional secondary products such as extract derivatives are expected to further amplify economic expansion (Moher et al., 2020). The need to maintain product consistency, while supporting innovation and development (Burgel et al., 2020) requires a better understanding of the physiological responses of cannabis to external stimuli. Research initiatives are needed to optimize current production strategies, enhancing our recognition of, and the precision at which we can invoke specific physiological responses to fit an assortment of industrial applications. Micropropagation offers unique opportunities to produce and maintain extensive populations of genetically uniform plantlets in time and cost-effective systems (Nathiya et al., 2013). Tissue culture techniques can be applied to examine essential plant responses to external stimuli in highly controlled environments under axenic conditions for biotechnological (Shukla et al., 2017), conservation (Ayuso et al., 2019) and various –omics related technologies (Andre et al., 2016). These approaches can be re-applied to suit the needs of the emerging cannabis industry.

Until recently, cannabis micropropagation has largely been an underground effort with few peer reviewed studies. This lack of insight concerning *in vitro* cannabis techniques has limited biotechnological utility of this crop (Smýkalová et al., 2019). While the current cannabis boom has led to the emergence of numerous *in vitro* protocols (Galán-Ávila et al., 2020; Lata et al., 2016; Wróbel et al., 2020), a robust and efficient protocol has yet to be fully developed. Several intrinsic (e.g., genotype, type, and age of explant) and extrinsic (e.g., basal salt medium, vitamins, plant growth regulators (PGRs), gelling agents, carbohydrate source, additional additives, temperature, and light) factors (Fig. 1) influence *in vitro* shoot growth and development and contribute to challenges in reproducibility (Hesami and Jones, 2020). Most previous studies in cannabis have investigated the effect of basal media, along with different types and concentrations of PGRs for shoot growth and regeneration (Chaohua et al., 2016; Lata et al., 2016; Movahedi and Torabi, 2015). Clonal line proliferation using apical and nodal explants on medium with reduced PGRs has been demonstrated as an effective approach for non-medicinal cannabis, while reducing the amount of emergent genetic variability (Wróbel et al., 2020). A cannabis micropropagation approach truly optimized for cross-cultivar maintenance and proliferation should allow formative physiological development on a pathway to photoautotrophic competence, coaxed through abiotic conditioning in the absence of PGRs. Though certain *in vitro* propagation, embryogenesis, and regeneration procedures rely on PGRs, many beneficial physiological characteristics can be induced or enhanced by appropriately adjusting light quality, quantity, and carbohydrate supply. *In vitro* shoot growth and development may be achieved or enhanced by manipulating light and sugar in the absence of PGRs.

**Figure 1.**
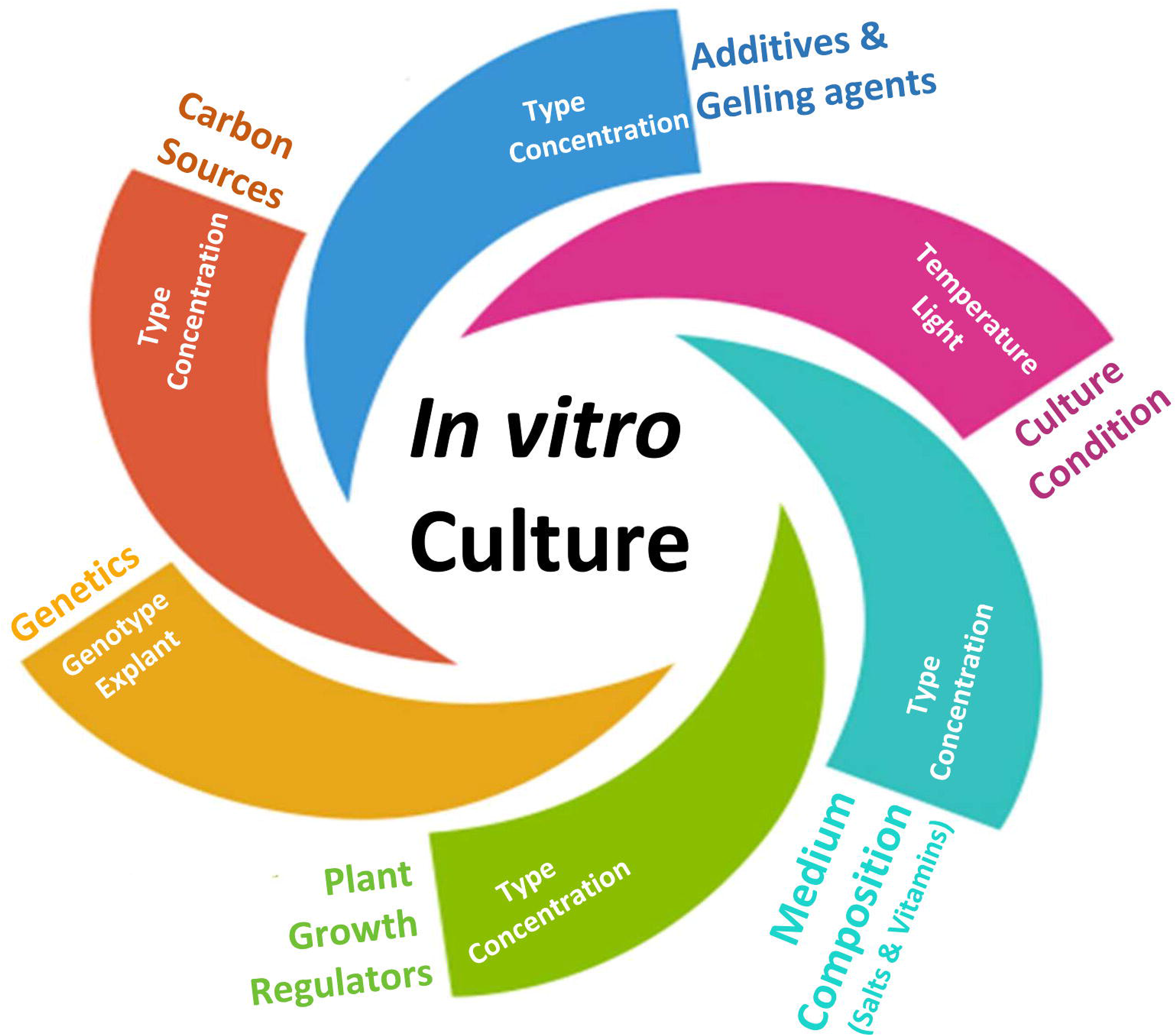
A schematic representation of factors influencing *in vitro* culture.

Though micropropagation protocols show promise to advance certain aspects of the cannabis industry, there remain issues with conventional *in vitro* systems. Photosynthetically incompetent organs, and fragile roots are phenotypic traits commonly observed in cultures (Jha and Bansal, 2012). Anatomical variations tend to be culture-induced, emerging with high humidity, elevated PGR concentrations, low light intensity, and high substrate water potential, causing physiological disorders such as hyperhydricity (Majada et al., 2000). These abiotic factors, along with limited culture CO_2_ availability, and potential for ethylene accumulation frequently impede photosynthetic responses (Nguyen et al., 2001), complicating the *ex vitro* transfer of specimens (Nathiya et al., 2013). Due to such limitations, new cultures must be supplied alternative carbon sources to maintain metabolic activity in otherwise daunting, closed environments (Eckstein et al., 2012). Sugar occurrence allows continuous plantlet development under low irradiance (Cioć et al., 2018), commonly used *in vitro*. A standardized addition of 3% (w/v) sucrose to the micropropagation media helps to counteract short-term, negative environmental impacts by providing substitute carbohydrates to elicit photo-mixotrophic metabolism (Gago et al., 2014). *In vitro* sucrose levels impact plantlet physiology by regulating genes relating to primary and secondary metabolic function (Yang et al., 2012). While supplemental carbon supply is a necessity for early-stage explants, developing plantlets can build sucrose dependence (Lembrechts et al., 2015), further limiting idealized physiological function and subsequent *ex vitro* re-localization. Conversely, previous work has also demonstrated that sugar can have positive effects on plantlet development under different environmental conditions *in vitro* (Eckstein et al., 2012; Kozai et al., 1987; Roh and Choi, 2004).

Although the occurrence of sucrose often activates photomixotrophic metabolic responses, light nevertheless bears high influence over *in vitro* success (Miler et al., 2019). Sugar and light signal essential metabolic processes which govern the condition of cultured plantlets (Eckstein et al., 2012). Though low light intensity *in vitro* hampers photosynthetic efficiency, overly high intensities can limit synthesis of photo-absorptive pigments and damage certain components of the photosynthetic apparatus (Cioć et al., 2018). Since high light levels throughout different culture stages can be stressful to developing plantlets, substitute carbon sources can help elicit photo-protective responses, indicating a possible sugar/light signalling pathway for photo-protection (Eckstein et al., 2012). Thus, photosynthetic limitation *in vitro* could largely be more related sub-optimal abiotic conditions in the presence of exogenous sugar, rather than the impact of the sugar itself (Arigita et al., 2002). Chloroplast localization (Eckstein et al., 2012), leaf area index (Snowden et al., 2016), and leaf thickness are influenced by changes in light quantity and quality. Proper development of these traits can increase photoabsorption saturation point (Macedo et al., 2011), enhancing plantlet fitness. Sustainable adjustment of the abiotic conditions combined with exogenous sugar can improve protective and repair responses (Eckstein et al., 2012; Tichá et al., 1998), allowing plantlets to more effectively sequester and utilize otherwise excessive and damaging photo-irradiation. Preliminary work conducted by our lab points in this direction in the case on micropropagated cannabis. Modifying abiotic factors and their interactions with sugar-related dynamics, is sometimes overlooked in micropropagation (Eckstein et al., 2012). Thus, research surrounding the potential to improve tissue culture protocols by optimizing abiotic influence and sugar-related dynamics should be thoroughly pursued.

The use of light emitting diodes (LEDs) for plant tissue culture allows strategic manipulation of light quality and intensity, impacting biomass and secondary metabolite accumulation of various species (Cioć et al., 2018; Manivannan et al., 2015; Ucar et al., 2016). Cool fluorescent lights have been popular for conventional micropropagation systems (Fanga et al., 2011) due to relatively low energy consumption, heat dissipation, and cost. However, they deliver light at wavelengths outside of the photoabsorptive range and lack control over spectral quality, which limits its power over physiological conditioning (Bello-Bello et al., 2016). There exists an established dogma that blue light (B) heavily influences chloroplast development, chlorophyll production, and stomata functionality, while red light (R) influences carbohydrate localization, and various anatomical processes such as leaf expansion (Hung et al., 2016; Ucar et al., 2016). Various combinations of these wavelengths can mutually and individually persuade shoot and root elongation (Ramírez-Mosqueda et al., 2017). LED technologies hold significant potential in the pursuit of plant growth in controlled environments, including plant tissue culture (Fontana et al., 2019). Control over spectral composition with LEDs allows wavelength emission that match photoreceptor action spectra to more directly trigger morphogenic responses (Li et al., 2010), while limiting heat dissipation, and energy consumption (Zhao et al., 2020). Photomorphogenic responses are primarily prompted by light quality through phytochrome reception of R and far-red light (Fr), and cryptochrome absorption of B (Miler and Zalewska, 2006), which largely shape plant development and physiology (Legris et al., 2019).

Despite the apparent simplicity of light quality and intensity, it is a complex factor comprised of nearly infinite potential mixtures which interact with other factors such as sucrose levels to influence *in vitro* shoot growth and development as a nonlinear, multifactorial, and complex process. The establishment and optimization of *in vitro* culture protocols have been principal challenges for many tissue culture researchers. Historically, micropropagation systems have been developed through serial manipulation and optimization of single factors, individually. Conventional statistical methods such as simple regression and ANOVA have typically been recommended for small databases with limited dimensions, and are therefore inappropriate for analyzing data derived from complex and non-linear processes such as light quality (Hesami et al., 2021b; Yoosefzadeh-Najafabadi et al., 2021a). The high probability of overfitting is one of the main disadvantages of using conventional statistical methods (Jafari and Shahsavar, 2020; Yoosefzadeh-Najafabadi et al., 2021b). Using conventional statistical methods, some of the puzzle pieces of *in vitro* practices have been sequentially assembled. However, many factors in tissue culture systems remain unoptimized. To overcome such setbacks, different factors can be simultaneously optimized through precision *in vitro* culture techniques using machine learning methods (Fig. 2). In recent years, machine learning algorithms such as artificial neural networks (ANNs) and neuro-fuzzy logic have been successfully applied for modeling and predicting various *in vitro* culture systems such as shoot growth and development, callogenesis, somatic embryogenesis, androgenesis, secondary metabolite production, and rhizogenesis (Hesami and Jones, 2020; Niazian and Niedbała, 2020). However, in most plant tissue culture studies, individual models were employed, and the efficiency of different machine learning algorithms has not been compared (Hesami et al., 2021c).

**Figure 2.**
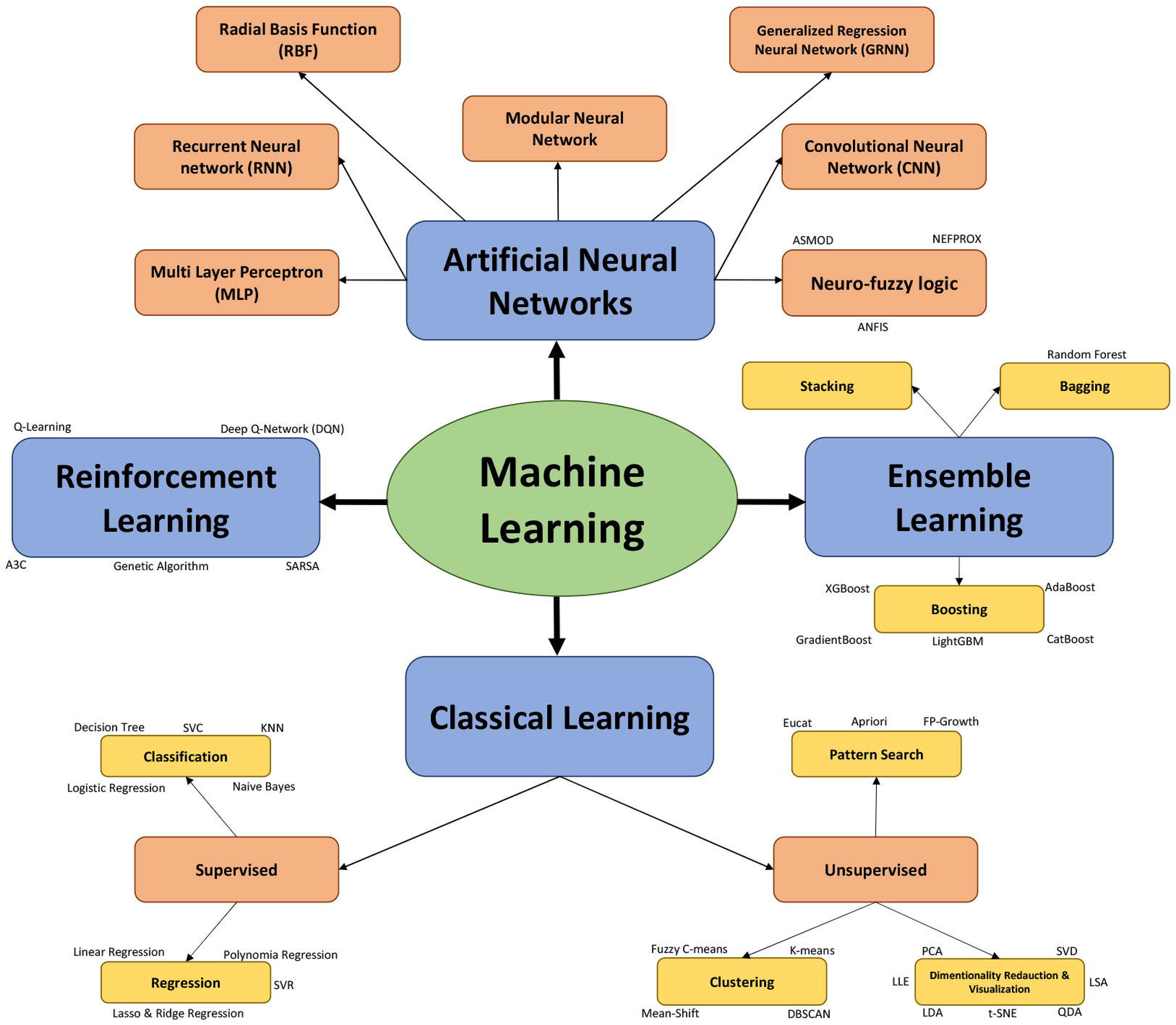
A schematic representation of different classes of machine learning algorithms.

There exist two general groups of optimization methods. Classical optimization algorithms include dynamic programming (DP), linear programming (LP), stochastic dynamic programming (SDP) which have limitations restricting their flexibility and efficiency. For instance, LP requires objective function and constraint to be linear, which is not ideal for plant tissue culture. Conversely, evolutionary optimization algorithms are considered more powerful mathematic methods for solving complex, multidimensional problems such as designating optimal factors for micropropagation with high accuracy and pace (Hesami and Jones, 2020). Although there are different types of evolutionary optimization algorithms, the genetic algorithm (GA) has been applied to the vast majority of plant tissue culture optimization studies relating to shoot proliferation, secondary metabolite production, and somatic embryogenesis. Despite the advantages that GA imparts over classical methods, premature convergence can sometimes lead to failure in obtaining a fully optimized solution (Hosseini-Moghari et al., 2015). To overcome this, new evolutionary optimization algorithms, including biogeography-based optimization (BBO), interior search algorithm (ISA), and symbiotic organisms search (SOS) have been developed. These approaches have been evaluated in different fields of study (Bozorg-Haddad et al., 2016; Hosseini-Moghari et al., 2015; Mokhtari Fard et al., 2012; Moravej and Hosseini-Moghari, 2016), and are expected to be superior in optimizing plant tissue culture protocols.

The current study tests the combined effects of B (400-500nm), R (600-700nm), Fr (700-800nm), and White (W) (400-700nm) (Figure 3, Figure 9) light at different intensities, and carbohydrate concentrations on shoot length, root length, number of nodes, number of shoots, and canopy surface area. Data collected were assessed using machine learning and evolutionary optimization algorithms to predict and optimize these factors for cannabis maintenance and proliferation *in vitro*. Predictions were then tested in a validation experiment to identify the best optimization algorithm for *in vitro* plant applications. Ultimately, the research presented will facilitate development of current practices for maintenance, proliferation, and acclimation of micropropagated cannabis, boosting our understanding of the dynamics between light and sugar-related plantlet responses, while identifying superior predictive analytic practices to guide future experimentation.

**Figure 3.**
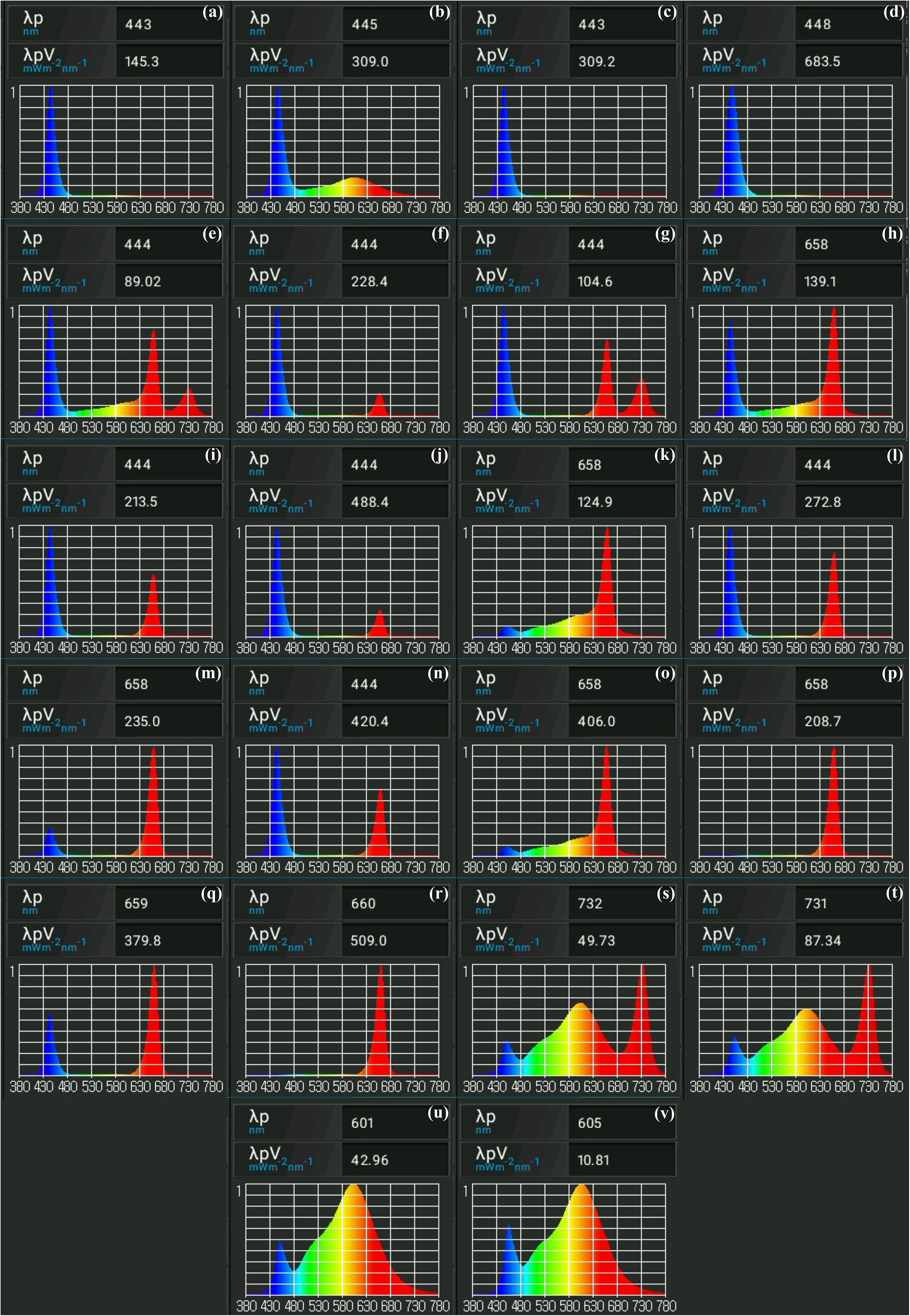
Spectral analyses of light treatments from the initial experiment. Images indicate relative amounts of fluencies emitted per treatment. Light spectra presented were obtained using Li-Cor LI-180 spectrometer. Presented are (a) 25 μmol/m^2^/s B + 25 μmol/m^2^/s W, (b) 50 μmol/m^2^/s B + 50 μmol/m^2^/s W, (c) 50 μmol/m^2^/s B, (d) 100 μmol/m^2^/s B, (e) 12.5 μmol/m^2^/s R + 12/5 μmol/m^2^/s B + 12.5 μmol/m^2^/s Fr + 12.5 μmol/m^2^/s W, (f) 12.5 μmol/m^2^/s R + 37.5 μmol/m^2^/s B, (g) 16.67 μmol/m^2^/s R + 16.67 μmol/m^2^/s B + 16.67 μmol/m^2^/s Fr, (h) 25 μmol/m^2^/s R + 25 μmol/m^2^/s B + 25 μmol/m^2^/s Fr + 25 μmol/m^2^/s W, (i) 25 μmol/m^2^/s R + 25 μmol/m^2^/s B, (j) 25 μmol/m^2^/s R + 75 μmol/m^2^/s B, (k) 25 μmol/m^2^/s R + 25 μmol/m^2^/s W, (l) 33.33 μmol/m^2^/s R + 33.33 μmol/m^2^/s B + 33.33 μmol/m^2^/s Fr, (m) 37.5 μmol/m^2^/s R + 12.5 μmol/m^2^/s B, (n) 50 μmol/m^2^/s R + 50 μmol/m^2^/s B, (o) 50 μmol/m^2^/s R + 50 μmol/m^2^/s W, (p) 50 μmol/m^2^/s R, (q) 75 μmol/m^2^/s R + 25 μmol/m^2^/s B, (r) 100 μmol/m^2^/s R, (s) 25 μmol/m^2^/s W + 25 μmol/m^2^/s Fr, (t) 50 μmol/m^2^/s W + 50 μmol/m^2^/s Fr, (u) 50 μmol/m^2^/s W, and (v) 100 μmol/m^2^/s W.

## 1 Materials and Methods

### 1.1 Plant material and experimental design

In this study, the effects of different light qualities, intensities and sucrose concentrations were evaluated for shoot growth, canopy surface area, and additional growth parameters, using the medicinal strain of cannabis “UP-802” supplied by Hexo, Brantford, ON. To this end, four plantlets per treatments were cultured in single Magenta boxes, allowing one experimental unit per treatment. Stock UP-802 specimens were maintained in cultures supplemented with 3% (w/v) sucrose, maintained under 16-hr photoperiod with 75% R, 12.5% B and 12.5% W LEDs at 50 μmol/m^2^/s. Both stock and experimental plantlets were grown at approximately 26◻. Non-experimental media components included 0.53% (w/v) DKW with vitamins, 0.10% (w/v) plant preservation mixture, and 0.60% (w/v) agar. Media pH was adjusted to 5.7 prior to agar addition, sterilization, and use. Chemicals were obtained from PhytoTech Labs.

To test the multivariable influences of sugar and light quality (intensity and spectrum) on *in vitro* cannabis development, plantlets were grown for 6-weeks with alternative sucrose concentrations, in compartmentalized light treatments. Programmable LED lights were used to provide light, allowing different combinations of B (400-500nm), R (600-700nm), Fr (700-800nm), and W (400-700nm) (Figure 3, Figure 9) light at specific intensities between 0-100 μmol/m2/s. The FinMax (bigfin.github.io/Prismatic) programmable LED lighting system was developed in-house to empower photobiology research with precise lighting treatments (Figure 3). The intensity of the 9 independently dimmable channels were programmed and calibrated at plant height using a spectrometer (Li-Cor LI-180).

At the end of each experiment, shoot length was measured by selecting the longest shoot and measuring from the root-shoot junction to apical meristem. Similarly, root length was measured from the root-shoot junction to root tip of the longest root. Number of nodes was collected by counting nodes on longest shoots. Shoot number was determined by counting emergent stems. Canopy surface areas were obtained by dissecting leaves and processing through ImageJ. All raw data were collected and processed using ImageJ software (Rueden et al., 2017).

For the preliminary experiment, apical explants were collected from stock UP-802 cultures, and sub-cultured to Magenta boxes with experimental media containing 1, 3 or 6% (w/v) sucrose. One culture of each sucrose concentration was randomly assigned to one of 22 different light treatments listed in Table 1, where they remained for 6-weeks. Data on shoot length, root length, number of shoots, number of nodes, and canopy surface area that was collected from 264 plantlets, as presented in Table 1, and processed with ImageJ software (Rueden et al., 2017). Raw experimental datasets were then analyzed using machine learning algorithms to build an appropriate model for cannabis shoot growth and development.

**Table 1.**
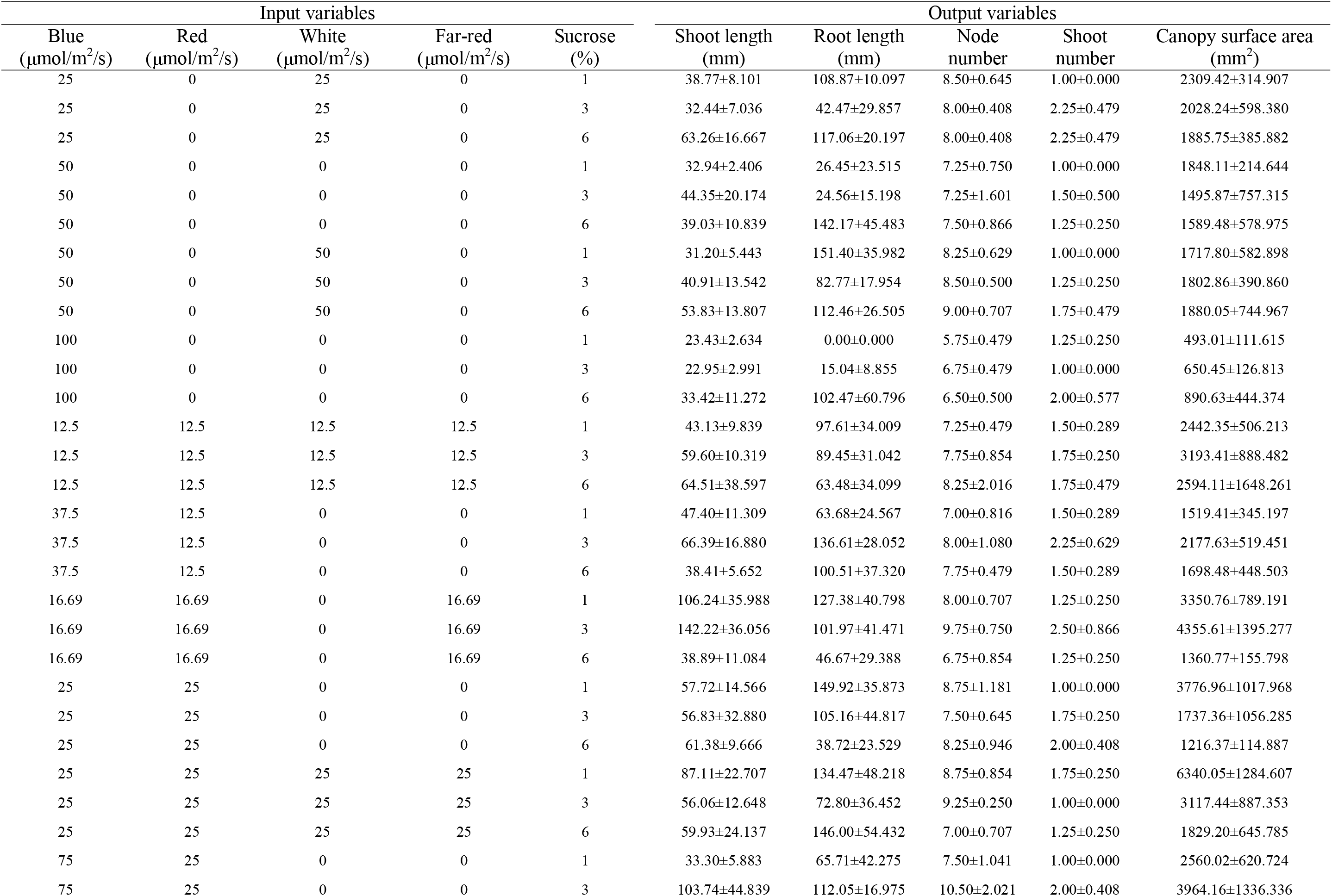
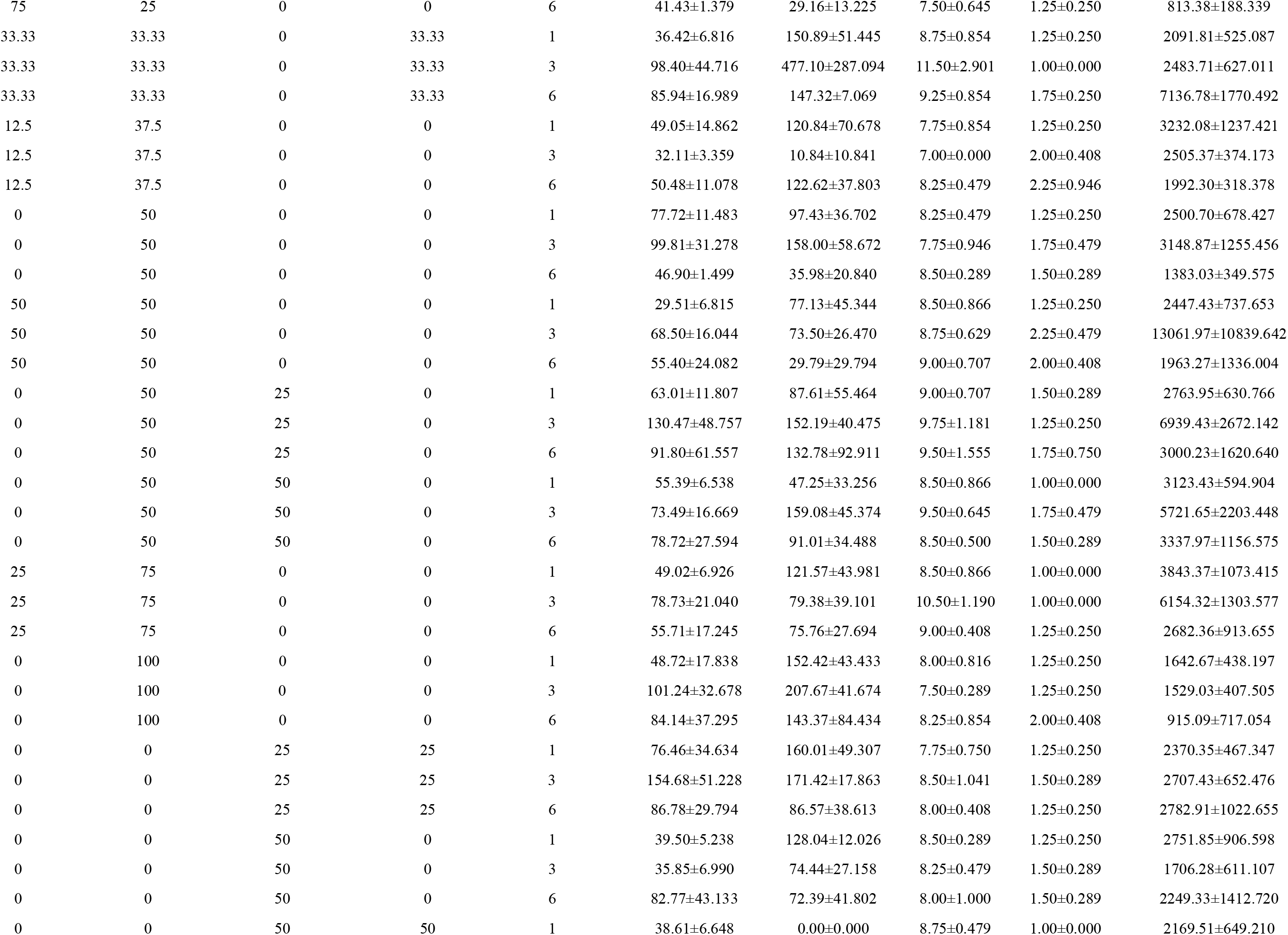
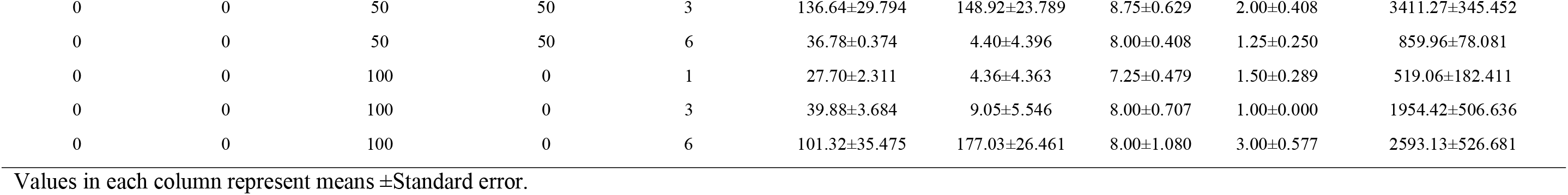
Effect of light and carbohydrate on *in vitro Cannabis* shoot growth and development.

| Input variables |  |  |  |  | Output variables |  |  |  |  |
| --- | --- | --- | --- | --- | --- | --- | --- | --- | --- |
| Blue<br>($\mu\text{mol}/\text{m}^2/\text{s}$) | Red<br>($\mu\text{mol}/\text{m}^2/\text{s}$) | White<br>($\mu\text{mol}/\text{m}^2/\text{s}$) | Far-red<br>($\mu\text{mol}/\text{m}^2/\text{s}$) | Sucrose<br>(%) | Shoot length<br>(mm) | Root length<br>(mm) | Node<br>number | Shoot<br>number | Canopy surface area<br>($\text{mm}^2$) |
| 25 | 0 | 25 | 0 | 1 | 38.77±8.101 | 108.87±10.097 | 8.50±0.645 | 1.00±0.000 | 2309.42±314.907 |
| 25 | 0 | 25 | 0 | 3 | 32.44±7.036 | 42.47±29.857 | 8.00±0.408 | 2.25±0.479 | 2028.24±598.380 |
| 25 | 0 | 25 | 0 | 6 | 63.26±16.667 | 117.06±20.197 | 8.00±0.408 | 2.25±0.479 | 1885.75±385.882 |
| 50 | 0 | 0 | 0 | 1 | 32.94±2.406 | 26.45±23.515 | 7.25±0.750 | 1.00±0.000 | 1848.11±214.644 |
| 50 | 0 | 0 | 0 | 3 | 44.35±20.174 | 24.56±15.198 | 7.25±1.601 | 1.50±0.500 | 1495.87±757.315 |
| 50 | 0 | 0 | 0 | 6 | 39.03±10.839 | 142.17±45.483 | 7.50±0.866 | 1.25±0.250 | 1589.48±578.975 |
| 50 | 0 | 50 | 0 | 1 | 31.20±5.443 | 151.40±35.982 | 8.25±0.629 | 1.00±0.000 | 1717.80±582.898 |
| 50 | 0 | 50 | 0 | 3 | 40.91±13.542 | 82.77±17.954 | 8.50±0.500 | 1.25±0.250 | 1802.86±390.860 |
| 50 | 0 | 50 | 0 | 6 | 53.83±13.807 | 112.46±26.505 | 9.00±0.707 | 1.75±0.479 | 1880.05±744.967 |
| 100 | 0 | 0 | 0 | 1 | 23.43±2.634 | 0.00±0.000 | 5.75±0.479 | 1.25±0.250 | 493.01±111.615 |
| 100 | 0 | 0 | 0 | 3 | 22.95±2.991 | 15.04±8.855 | 6.75±0.479 | 1.00±0.000 | 650.45±126.813 |
| 100 | 0 | 0 | 0 | 6 | 33.42±11.272 | 102.47±60.796 | 6.50±0.500 | 2.00±0.577 | 890.63±444.374 |
| 12.5 | 12.5 | 12.5 | 12.5 | 1 | 43.13±9.839 | 97.61±34.009 | 7.25±0.479 | 1.50±0.289 | 2442.35±506.213 |
| 12.5 | 12.5 | 12.5 | 12.5 | 3 | 59.60±10.319 | 89.45±31.042 | 7.75±0.854 | 1.75±0.250 | 3193.41±888.482 |
| 12.5 | 12.5 | 12.5 | 12.5 | 6 | 64.51±38.597 | 63.48±34.099 | 8.25±2.016 | 1.75±0.479 | 2594.11±1648.261 |
| 37.5 | 12.5 | 0 | 0 | 1 | 47.40±11.309 | 63.68±24.567 | 7.00±0.816 | 1.50±0.289 | 1519.41±345.197 |
| 37.5 | 12.5 | 0 | 0 | 3 | 66.39±16.880 | 136.61±28.052 | 8.00±1.080 | 2.25±0.629 | 2177.63±519.451 |
| 37.5 | 12.5 | 0 | 0 | 6 | 38.41±5.652 | 100.51±37.320 | 7.75±0.479 | 1.50±0.289 | 1698.48±448.503 |
| 16.69 | 16.69 | 0 | 16.69 | 1 | 106.24±35.988 | 127.38±40.798 | 8.00±0.707 | 1.25±0.250 | 3350.76±789.191 |
| 16.69 | 16.69 | 0 | 16.69 | 3 | 142.22±36.056 | 101.97±41.471 | 9.75±0.750 | 2.50±0.866 | 4355.61±1395.277 |
| 16.69 | 16.69 | 0 | 16.69 | 6 | 38.89±11.084 | 46.67±29.388 | 6.75±0.854 | 1.25±0.250 | 1360.77±155.798 |
| 25 | 25 | 0 | 0 | 1 | 57.72±14.566 | 149.92±35.873 | 8.75±1.181 | 1.00±0.000 | 3776.96±1017.968 |
| 25 | 25 | 0 | 0 | 3 | 56.83±32.880 | 105.16±44.817 | 7.50±0.645 | 1.75±0.250 | 1737.36±1056.285 |
| 25 | 25 | 0 | 0 | 6 | 61.38±9.666 | 38.72±23.529 | 8.25±0.946 | 2.00±0.408 | 1216.37±114.887 |
| 25 | 25 | 25 | 25 | 1 | 87.11±22.707 | 134.47±48.218 | 8.75±0.854 | 1.75±0.250 | 6340.05±1284.607 |
| 25 | 25 | 25 | 25 | 3 | 56.06±12.648 | 72.80±36.452 | 9.25±0.250 | 1.00±0.000 | 3117.44±887.353 |
| 25 | 25 | 25 | 25 | 6 | 59.93±24.137 | 146.00±54.432 | 7.00±0.707 | 1.25±0.250 | 1829.20±645.785 |
| 75 | 25 | 0 | 0 | 1 | 33.30±5.883 | 65.71±42.275 | 7.50±1.041 | 1.00±0.000 | 2560.02±620.724 |
| 75 | 25 | 0 | 0 | 3 | 103.74±44.839 | 112.05±16.975 | 10.50±2.021 | 2.00±0.408 | 3964.16±1336.336 |
| 75 | 25 | 0 | 0 | 6 | 41.43±1.379 | 29.16±13.225 | 7.50±0.645 | 1.25±0.250 | 813.38±188.339 |
| 33.33 | 33.33 | 0 | 33.33 | 1 | 36.42±6.816 | 150.89±51.445 | 8.75±0.854 | 1.25±0.250 | 2091.81±525.087 |
| 33.33 | 33.33 | 0 | 33.33 | 3 | 98.40±44.716 | 477.10±287.094 | 11.50±2.901 | 1.00±0.000 | 2483.71±627.011 |
| 33.33 | 33.33 | 0 | 33.33 | 6 | 85.94±16.989 | 147.32±7.069 | 9.25±0.854 | 1.75±0.250 | 7136.78±1770.492 |
| 12.5 | 37.5 | 0 | 0 | 1 | 49.05±14.862 | 120.84±70.678 | 7.75±0.854 | 1.25±0.250 | 3232.08±1237.421 |
| 12.5 | 37.5 | 0 | 0 | 3 | 32.11±3.359 | 10.84±10.841 | 7.00±0.000 | 2.00±0.408 | 2505.37±374.173 |
| 12.5 | 37.5 | 0 | 0 | 6 | 50.48±11.078 | 122.62±37.803 | 8.25±0.479 | 2.25±0.946 | 1992.30±318.378 |
| 0 | 50 | 0 | 0 | 1 | 77.72±11.483 | 97.43±36.702 | 8.25±0.479 | 1.25±0.250 | 2500.70±678.427 |
| 0 | 50 | 0 | 0 | 3 | 99.81±31.278 | 158.00±58.672 | 7.75±0.946 | 1.75±0.479 | 3148.87±1255.456 |
| 0 | 50 | 0 | 0 | 6 | 46.90±1.499 | 35.98±20.840 | 8.50±0.289 | 1.50±0.289 | 1383.03±349.575 |
| 50 | 50 | 0 | 0 | 1 | 29.51±6.815 | 77.13±45.344 | 8.50±0.866 | 1.25±0.250 | 2447.43±737.653 |
| 50 | 50 | 0 | 0 | 3 | 68.50±16.044 | 73.50±26.470 | 8.75±0.629 | 2.25±0.479 | 13061.97±10839.642 |
| 50 | 50 | 0 | 0 | 6 | 55.40±24.082 | 29.79±29.794 | 9.00±0.707 | 2.00±0.408 | 1963.27±1336.004 |
| 0 | 50 | 25 | 0 | 1 | 63.01±11.807 | 87.61±55.464 | 9.00±0.707 | 1.50±0.289 | 2763.95±630.766 |
| 0 | 50 | 25 | 0 | 3 | 130.47±48.757 | 152.19±40.475 | 9.75±1.181 | 1.25±0.250 | 6939.43±2672.142 |
| 0 | 50 | 25 | 0 | 6 | 91.80±61.557 | 132.78±92.911 | 9.50±1.555 | 1.75±0.750 | 3000.23±1620.640 |
| 0 | 50 | 50 | 0 | 1 | 55.39±6.538 | 47.25±33.256 | 8.50±0.866 | 1.00±0.000 | 3123.43±594.904 |
| 0 | 50 | 50 | 0 | 3 | 73.49±16.669 | 159.08±45.374 | 9.50±0.645 | 1.75±0.479 | 5721.65±2203.448 |
| 0 | 50 | 50 | 0 | 6 | 78.72±27.594 | 91.01±34.488 | 8.50±0.500 | 1.50±0.289 | 3337.97±1156.575 |
| 25 | 75 | 0 | 0 | 1 | 49.02±6.926 | 121.57±43.981 | 8.50±0.866 | 1.00±0.000 | 3843.37±1073.415 |
| 25 | 75 | 0 | 0 | 3 | 78.73±21.040 | 79.38±39.101 | 10.50±1.190 | 1.00±0.000 | 6154.32±1303.577 |
| 25 | 75 | 0 | 0 | 6 | 55.71±17.245 | 75.76±27.694 | 9.00±0.408 | 1.25±0.250 | 2682.36±913.655 |
| 0 | 100 | 0 | 0 | 1 | 48.72±17.838 | 152.42±43.433 | 8.00±0.816 | 1.25±0.250 | 1642.67±438.197 |
| 0 | 100 | 0 | 0 | 3 | 101.24±32.678 | 207.67±41.674 | 7.50±0.289 | 1.25±0.250 | 1529.03±407.505 |
| 0 | 100 | 0 | 0 | 6 | 84.14±37.295 | 143.37±84.434 | 8.25±0.854 | 2.00±0.408 | 915.09±717.054 |
| 0 | 0 | 25 | 25 | 1 | 76.46±34.634 | 160.01±49.307 | 7.75±0.750 | 1.25±0.250 | 2370.35±467.347 |
| 0 | 0 | 25 | 25 | 3 | 154.68±51.228 | 171.42±17.863 | 8.50±1.041 | 1.50±0.289 | 2707.43±652.476 |
| 0 | 0 | 25 | 25 | 6 | 86.78±29.794 | 86.57±38.613 | 8.00±0.408 | 1.25±0.250 | 2782.91±1022.655 |
| 0 | 0 | 50 | 0 | 1 | 39.50±5.238 | 128.04±12.026 | 8.50±0.289 | 1.25±0.250 | 2751.85±906.598 |
| 0 | 0 | 50 | 0 | 3 | 35.85±6.990 | 74.44±27.158 | 8.25±0.479 | 1.50±0.289 | 1706.28±611.107 |
| 0 | 0 | 50 | 0 | 6 | 82.77±43.133 | 72.39±41.802 | 8.00±1.000 | 1.50±0.289 | 2249.33±1412.720 |
| 0 | 0 | 50 | 50 | 1 | 38.61±6.648 | 0.00±0.000 | 8.75±0.479 | 1.00±0.000 | 2169.51±649.210 |
| 0 | 0 | 50 | 50 | 3 | 136.64±29.794 | 148.92±23.789 | 8.75±0.629 | 2.00±0.408 | 3411.27±345.452 |
| 0 | 0 | 50 | 50 | 6 | 36.78±0.374 | 4.40±4.396 | 8.00±0.408 | 1.25±0.250 | 859.96±78.081 |
| 0 | 0 | 100 | 0 | 1 | 27.70±2.311 | 4.36±4.363 | 7.25±0.479 | 1.50±0.289 | 519.06±182.411 |
| 0 | 0 | 100 | 0 | 3 | 39.88±3.684 | 9.05±5.546 | 8.00±0.707 | 1.00±0.000 | 1954.42±506.636 |
| 0 | 0 | 100 | 0 | 6 | 101.32±35.475 | 177.03±26.461 | 8.00±1.080 | 3.00±0.577 | 2593.13±526.681 |

Values in each column represent means ±Standard error.

### 1.2 Modeling procedure

Three well-known machine learning algorithms, MLP, GRNN, and ANFIS, were applied to model and predict *in vitro* shoot growth and development of cannabis using the collected dataset. Box-Cox transformation was employed to normalize the data before using the machine learning algorithms. Principal component analysis (PCA) was applied to detect outliers, but no outliers were identified. In this study, the five-fold cross-validation approach, with 10 repetitions was applied to evaluate the prediction accuracy of the tested machine learning algorithms.

Different light qualities (B, R, W, and Fr) at various intensities and different levels of Sucrose were selected as input variables, while shoot length, root length, number of nodes, number of shoots, and canopy surface area were considered as target (output) variables (Fig. 4a).

**Figure 4.**
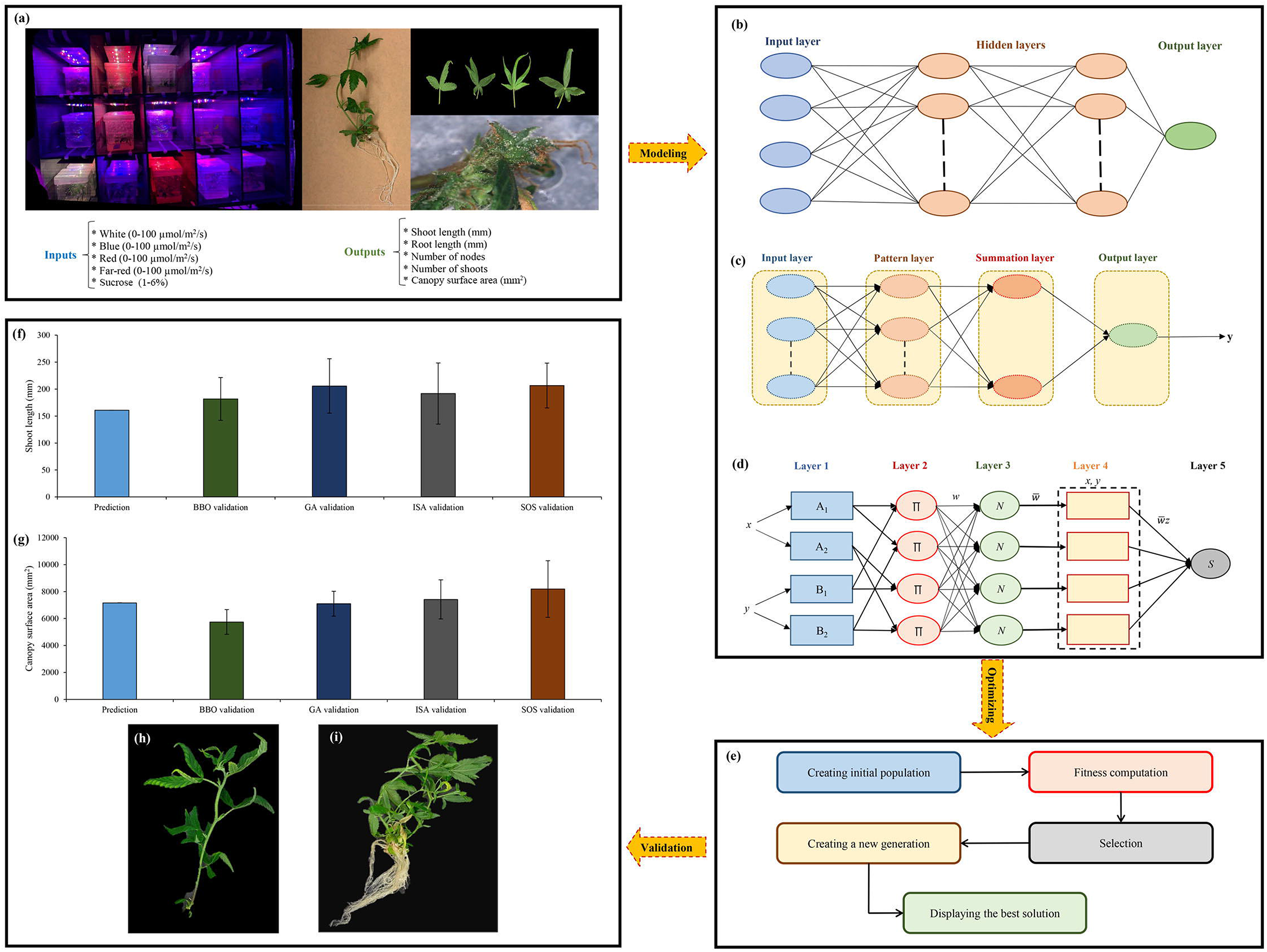
Step-by-step methodology of the current study, including (a) data obtained, (b-d) data modeling through multilayer perceptron (MLP), generalized regression neural networks (GRNN), and adaptive neuro-fuzzy inference system (ANFIS), respectively, (e) main steps of optimization process through different optimization algorithms, (f,g) results of the validation experiment for shoot growth and canopy surface area, respectively, and (h,i) shoot growth and canopy surface area obtained from symbiotic organisms search (SOS).

To evaluate and compare the efficiency and accuracy of the machine learning algorithms, R^2^ (coefficient of determination), mean bias error (MBE), and root mean square error (RMSE) were employed based on the following equations:

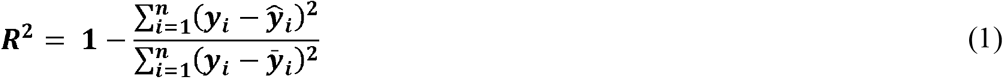

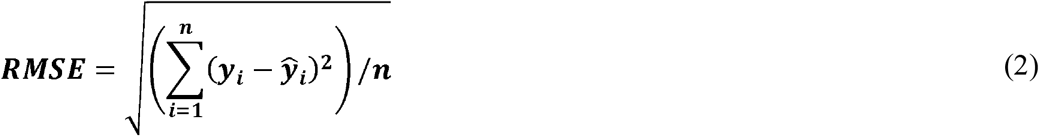

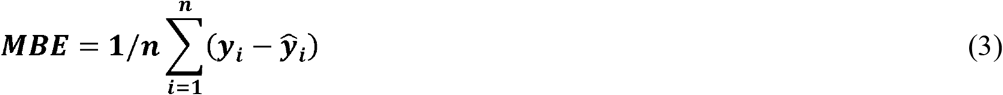

Where *y*_*i*_ is the value of prediction, *n* is the number of data, and 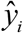 is value of observation.

#### 2.2.1 Multi-Layer Perceptron (MLP)

MLP belongs to the ANNs which is inspired by the neural structure of the human brain. A neuron in the human neural network receives impulses by using a number of dendrites from other neurons. Based on the received impulses, a neuron through its single axon may send a signal to other neurons. Like the human neural network, ANNs contain nodes, each of which receives a number of input variables and produce a single target variable, where the target variable is a relatively simple function of the input variables (Fig. 4b).

The 3-layer backpropagation MLP is a parallel and distributed algorithm that uses supervised learning for the training subset. The following equation is employed to minimize the error between the input and target variables:

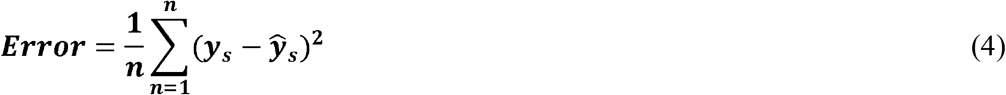

Where *y*_*s*_ is the s^th^ observed variable, *n* is the number of observations, and 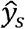 is the s^th^ predicted variable.

To determine the 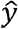 in the model *k* output variables and with *p* neurons in the hidden layer, following function is employed:

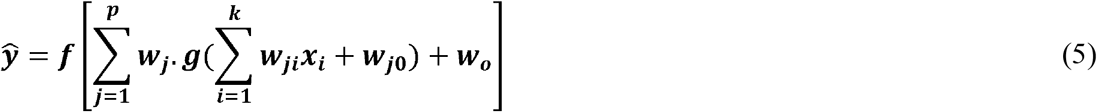

where *w*_*j*_ represents the weighted input data into the j^th^ neuron of the hidden layer, *w*_*0*_ equals the bias connected to the neuron of output, *w*_*ji*_ represents the weight of the direct relationship of input neuron *i* to the hidden neuron *j*, *x*_*i*_ is the i^th^ target variable, *f* represents activation function for the target neuron, *w*_*j0*_ shows the bias for node j^th^, and *g* shows the activation function for the hidden neuron.

Since the number of hidden units and the number of neurons in each node play an important role in the efficiency of MLP, they should be determined. In the present investigation, trial and error-based approach was used to detect the optimal neuron number in the hidden layer. Also, linear function (purelin) as the transfer functions of output layer and hyperbolic tangent sigmoid function (tansig) as the transfer functions of hidden layer were applied. Moreover, A Levenberg-Marquardt algorithm was employed for adjusting bias and weights.

#### 2.2.2 Generalized Regression Neural Network (GRNN)

The GRNN as another kind of ANNs consists of four layers (Fig. 4c). The node in input layer completely enters the node in pattern layer. The output of each neuron in pattern layer is connected to the summation neurons. The unweighted pattern neuron outputs are determined by D-summation

neuron, while the weighted pattern neuron outputs are computed by S-summation neuron. Finally, the following equation is employed to determine the output:

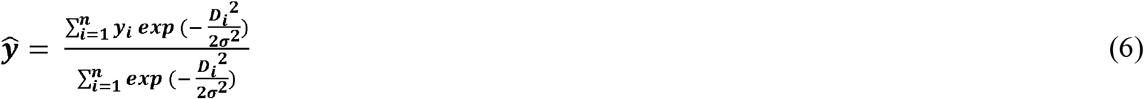

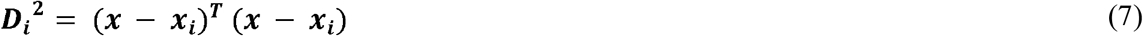

where σ represents width parameter, 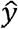 shows the average of all the weighted observed output data, *y*_*i*_ shows the i^th^ output variable, and *D*_*i*_^2^ equals a scalar function which is based on any *x*_*i*_ and *y*_*i*_ observed data.

### 2.2.3 Adaptive Neuro-Fuzzy Inference System (ANFIS)

ANFIS developed by Jang (1993) is one of the most well-known neuro-fuzzy logic systems. The overall ANFIS model with two Takagi and Sugeno type if-then rules can be defined as follow:

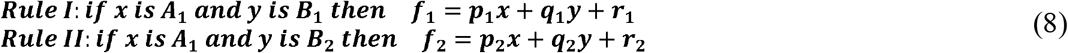

Where *x* and *y* are input variables; *f*_*1*_ and *f*_*2*_ are the outputs within the fuzzy area determined by the fuzzy rule; *A*_*1*_, *A*_*2*_, *B*_*1*_, and *B*_*2*_ are the fuzzy sets; *p*_*1*_, *p*_*2*_, *q*_*1*_, *q*_*2*_, *r*_*1*_, and *r*_*2*_ are the design parameters that are specified during the training set. The ANFIS model is built of five layers (Fig. 4d) as follow:

Layer 1 (adaptive or input layer): Every adaptive (input) node *i* in layer 1 defines a square node with a node function:

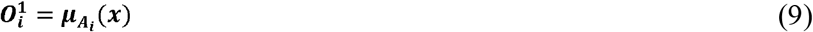

Where 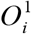 is the fuzzy membership grade, x is the input of adaptive node *i*, and 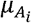 is Gaussian membership function which is deremined as follow:

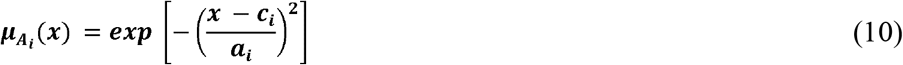

where *a*_*i*_ and *c*_*i*_ are premise parameters.

Layer 2 (rule layer): Every role node in layer 2 can be considered as a circle node labeled ∏ where the output is the result of all incoming inputs.

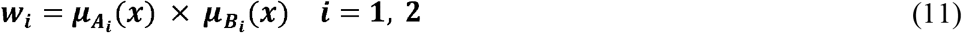

Each node output displays the rule’s firing strength.

Layer 3 (average layer): Each node as a fixed node in layer 3 is labeled *N*. The *i*^*th*^ node determines the ratio of the firing strength of *i*^*th*^ rule to the total rules’ firing strengths. The outputs of layer 3 (normalized firing strengths) are calculated as follow:

Where 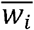 is output of this layer.

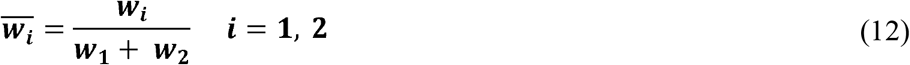

Layer 4 (consequent layer): Nodes in layer 4 are called consequent nodes. The following equation is used to calculate the output of this layer.

Where *p*_*i*_, *q*_*i*_, and *r*_*i*_ are parameter sets and 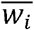 is output of layer 3.

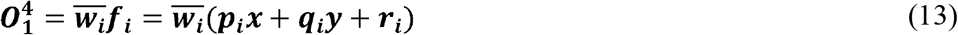

Layer 5 (output layer): There is only one single fixed node labeled *S* in this layer. The final output 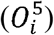 of the model is calculated based on the following equation:

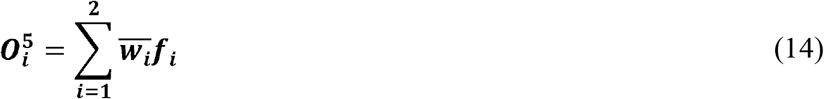

In the current study, the Gaussian membership function (between 3 and 5 membership functions for different variables) was considered based on a trial and error approach. The number of epochs to train the models was also set to 10. Moreover, the least-squares method and backpropagation algorithm were applied to adjust the consequent and premise parameters, respectively.

### 1.3 Sensitivity analysis

Sensitivity analysis was performed to assess the degree of importance of various forms of light (B, R, W, and Fr) and exogenous carbohydrates on shoot length, root length, number of nodes, number of shoots, and canopy surface area by determining the variable sensitivity ratio (VSR). VSR can be defined as the ratio of variable sensitivity error (VSE) to the RMSE of the developed model. A greater VSR shows a higher degree of importance.

### 1.4 Optimization procedure

In the current study, four different single-objective evolutionary optimization algorithms including BBO, ISA, SOS, and GA were separately employed to find optimal levels of input variables (Sucrose, B, R, W, and Fr) for maximizing each fitness function (shoot length, root length, number of nodes, number of shoots, and canopy surface area). Generally, evolutionary optimization algorithms consist of five main steps including creating an initial population, fitness computation, selection, creating a new generation, and displaying the best solution (Fig. 4e). The details of each algorithm have been presented below.

#### 2.4.1 Biogeography-Based Optimization (BBO)

The term “Biogeography” refers to the study of ecosystems and the geographical distribution of species. BBO introduced by Simon (2008) is based on biogeographic concepts such as migration, evolution, adaptation, and extinction of organisms among habitats. In theory, appropriate regions for living organism’s settlement are defined by the habitat suitability index (HSI) that depends on several factors such as precipitation, temperature, area, and vegetative cover which are known as suitability index variables (SIVs). Indeed, HIS as a dependent variable is determined by SIVs as independent variables. Therefore, more living organisms can be accommodated in habitats with higher values of HIS and vice versa, lower HSI values support fewer organisms. Subsequently, a stronger tendency for living organisms to emigrate from the habitat to find new places with lower population density and more suitable conditions can be seen by increasing the number of species in a habitat.

The highest λ can be seen when there are no species in the habitat. The λ decreases by increasing the number of species in the habitat and, finally, the λ becomes zero when the habitat capacity is completed (the maximum number of species in the habitat equals *S*_*max*_). On the other hand, the μ enhances by increasing the number of species in the habitat until the habitat becomes empty. Hence, the equilibrium number of species in the habitat can be seen when λ equals μ. Generally, λ and μ can be determined based on the following equations:

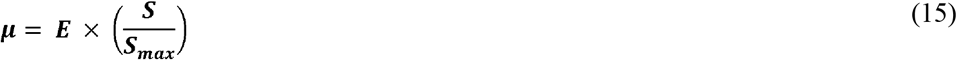

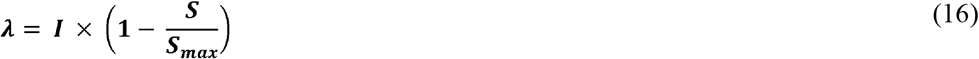

Where *S* is the number of species, *I* shows the maximum rate of immigration, and *E* is the maximum rate of emigration.

In the BBO method, habitat and SIVs play the role of solution and the decision variables, respectively. Therefore, the HSI can be considered as the objective function in this optimization algorithm. If there is a particular graph with *E*=*I* for each solution, HSI has a direct relationship with *S*, in which case HSI values can be used instead of *S*. The step-by-step procedure of the ISA method has been presented in Figure 5.

**Figure 5.**
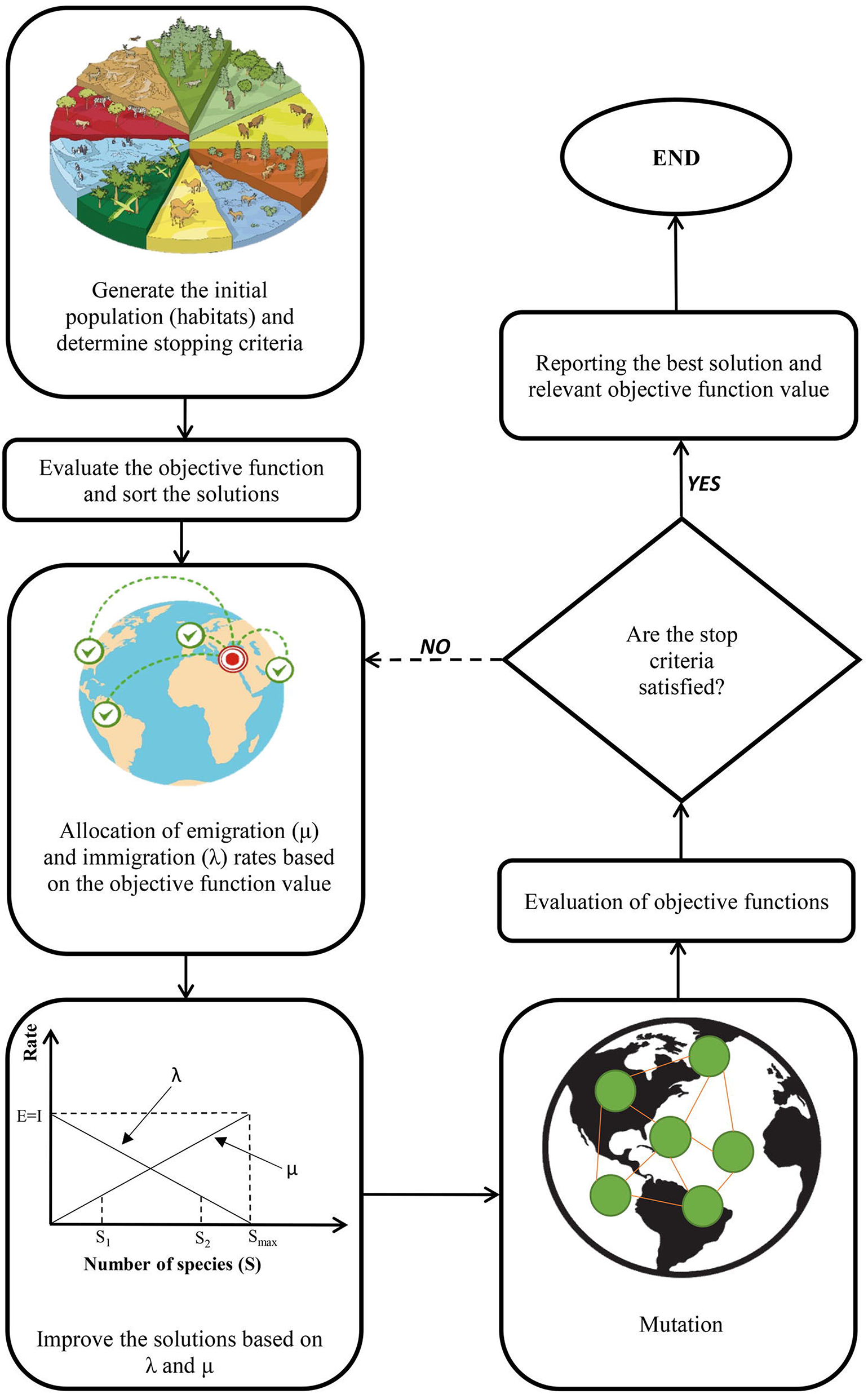
A schematic representation of biogeography-based optimization (BBO) algorithm.

With a specific probability *P*_*mod*_, different solutions can help each other for improvement. If the *S*_*i*_ is selected as an improvement, the λ is employed to adjust its SIVs. Subsequently, the μ relevant to other solutions is applied to choose the improved solution. The SIVs of the *S*_*i*_ solution are then used for randomly replacing SIVs from selected solutions. The suitable values of μ can be arbitrarily considered by using an arithmetic progression, with the common difference of successive members equal to 1/(population size −1), between 0 and 1. After calculation of μ, λ can be determined as λ=1−μ.

For lack of elitism, all solutions should be modified at all steps. However, modifying the amount of any solution is conversely related to its HSI. A roulette wheel is used for choosing the modifier solution which is based on a probability proportional to the μ. Transferring SIVs, as an inferior strategy, from one solution to another solution restricts the search choices within the decision space. Therefore, the following equation has been recommended for replacing SIVs:

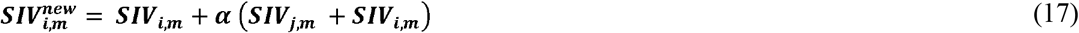

Where 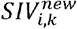 equals *m*^*th*^ modified SIV of the *i*^*th*^ solution, *α* is a parameter between 0 and 1, which is determined by the user, *SIV*_*j,m*_ is *m*^*th*^ SIV of the *j*^*th*^ solution, *SIV*_*j,m*_ equals *m*^*th*^ SIV of the *i*^*th*^ solution.

Severe catastrophes such as natural hazards, the spreading of infectious diseases, and other catastrophes can quickly change the HSI of a habitat. These unfavorable conditions act like mutations in GA.

#### 2.4.2 Interior Search Algorithm (ISA)

The ISA method introduced by Gandomi, (2014) is based on the concepts of interior design and decoration using mirrors, such that, several mirrors can be used to create a more decorative environment. To meet decoration project goals, it is necessary to satisfy the desires of the clients’ desires using available resources. The interior design commences with centering bounded elements to create a more appealing interior vista based on client approval. The ISA method is inspired by this repetitive process to solve optimization problems. With this algorithm, an element can only be moved to a position allowing a more decorative view (better fitness) while satisfying customer resource and need demands (constraints).

The most important step of interior design is positioning the mirrors by the fittest and most striking elements to highlighting their attractiveness. Generally, the elements are classified in two ways (i) the composition category, which is applied for composition optimization, and (ii) the mirror category, which is employed for mirror search. Therefore, the ISA method can be explained as follow.

1. Create the position of elements between upper bound (UB) and lower bound (LB) randomly and determine their fitness value.
2. Discover the element with minimum objective function in minimization problem (the fittest element) in *j*^*th*^ iteration.
3. Apply a random variable *r*1 (ranging between 0 and1 for each element) and *α* as a threshold value (*α* is also a value between 0 and 1) to divide other elements, except the fittest element, into mirror category and composition category. Elements with *α* ≤◻*r*1 go to the composition category; otherwise, they go to the mirror category. Since *a* is the only parameter of the ISA method, it is necessary to carefully tune *α* for obtaining balance between diversification and intensification. In the current study, a linear equation from 0.1 to 0.9 was used for determining the value of *α* during optimization iterations, meaning the *α* value modifies as iteration goes up towards its maximum number. This method provides a parametric optimization algorithm in which the algorithm can automatically adjust its parameter. As the iteration approaches the highest iteration number, the *α* value reaches 0.9. Subsequently, the optimization procedure slowly shifts to mirror search to promote exploitation at the end of repetitions.
4. For the fittest element, it is beneficial to lightly change positions using the random walk for a local search around the fittest element. The following equation can be used for calculating the fittest element.

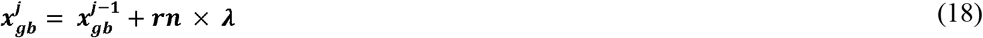 Where, 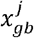 is the fittest element, λ is scale factor = 0.01(UB◻−◻LB), and *rn* presents vector of normally distributed random numbers.
5. For the composition category, each element in this category is randomly displaced. The following equation is used for determining the changes in UB and LB:

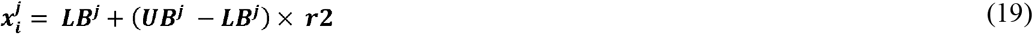 Where 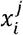 shows *i*^*th*^ element in the *j*^*th*^ iteration, UB^j^ and *LB*^*j*^ represent upper and lower bounds of the class in *j*^*th*^ iteration, respectively, and *r*2 is random value between 0 and 1.
6. For the mirror category, a mirror is randomly placed between the fittest element and each composition element. The following equation is applied for calculating the position of a mirror for the *i*^*th*^ element of the *j*^*th*^ iteration:

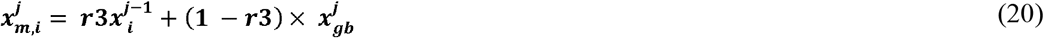 Where 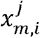 equals the position of a mirror for the *i*^*th*^ element of the *j*^*th*^ iteration, and *r*3 is a random value between 0 and 1. The virtual location of the element (the image of the element in the mirror) depends on the position of the mirror and is calculated based on the following equation:

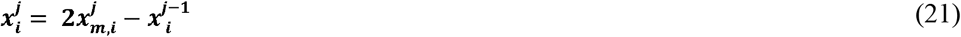
7. The virtual elements and fitness values of the new positions of the elements should be determined. The positions should be updated if their finesses are improved. It can be calculated based on the following equation (for a minimization problem):

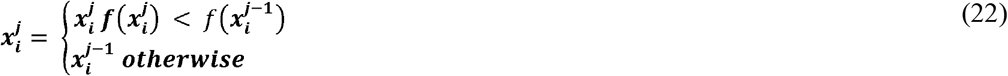
8. If any of the termination criteria are not satisfied, the steps should be repeated from step 2. The step-by-step procedure of the ISA method has been presented in Figure 6.

**Figure 6.**
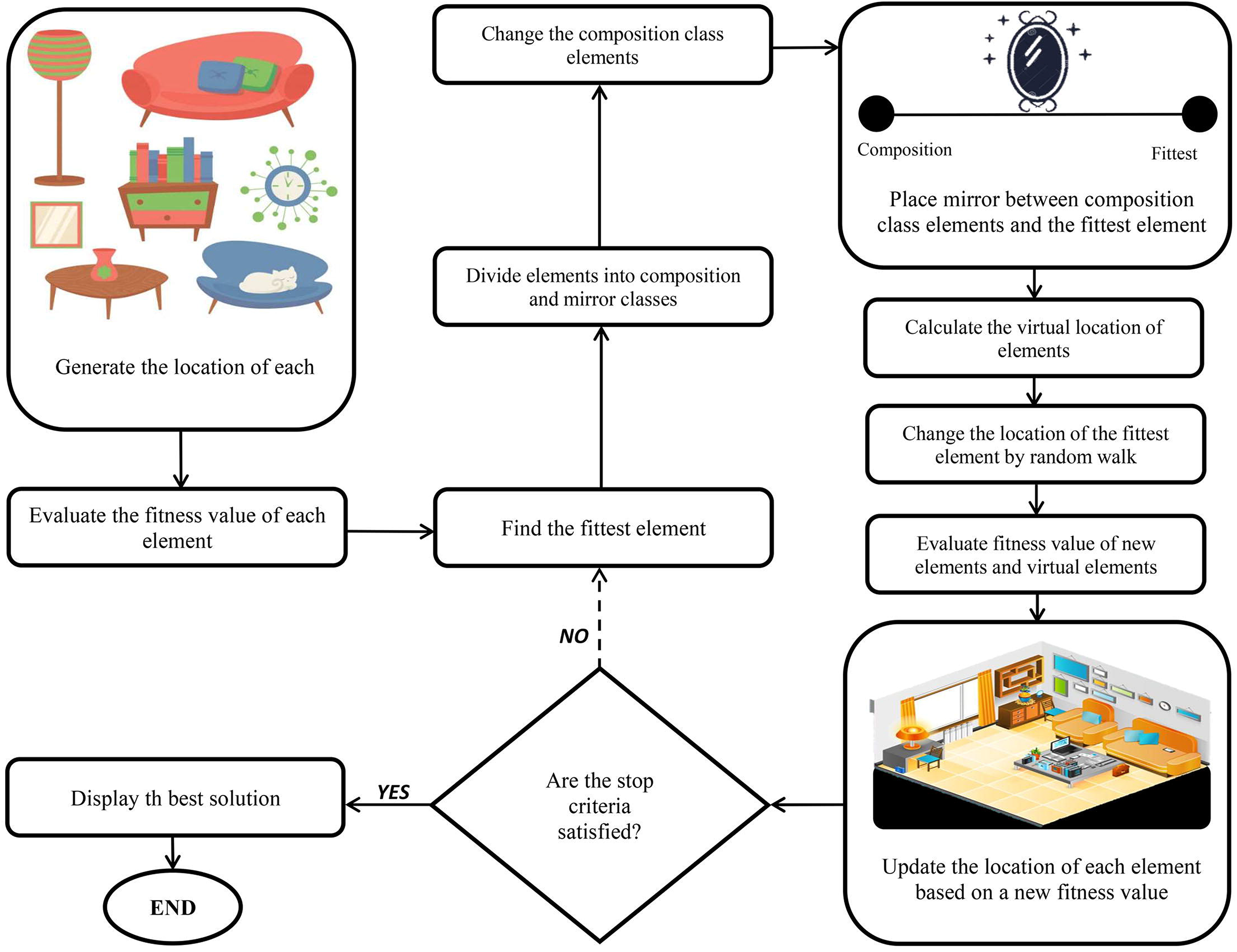
A schematic representation of interior search algorithm (ISA) algorithm.

#### 2.4.3 Symbiotic organisms search (SOS) algorithm

The SOS introduced by Cheng and Prayogo (2014) can be considered a nature-inspired optimization method. The SOS algorithm simulates three various interactions of symbioses amongst species of an ecosystem. Much like the majority of evolutionary optimization algorithms, SOS creates an ecosystem as an initial population plus particular operators through an iterative method to find a near-optimal solution among candidate organisms as possible solutions within the promising space of a search area. However, the SOS method does not reproduce offspring. Step-by-step SOS procedure methods are presented in Figure 7.

**Figure 7.**
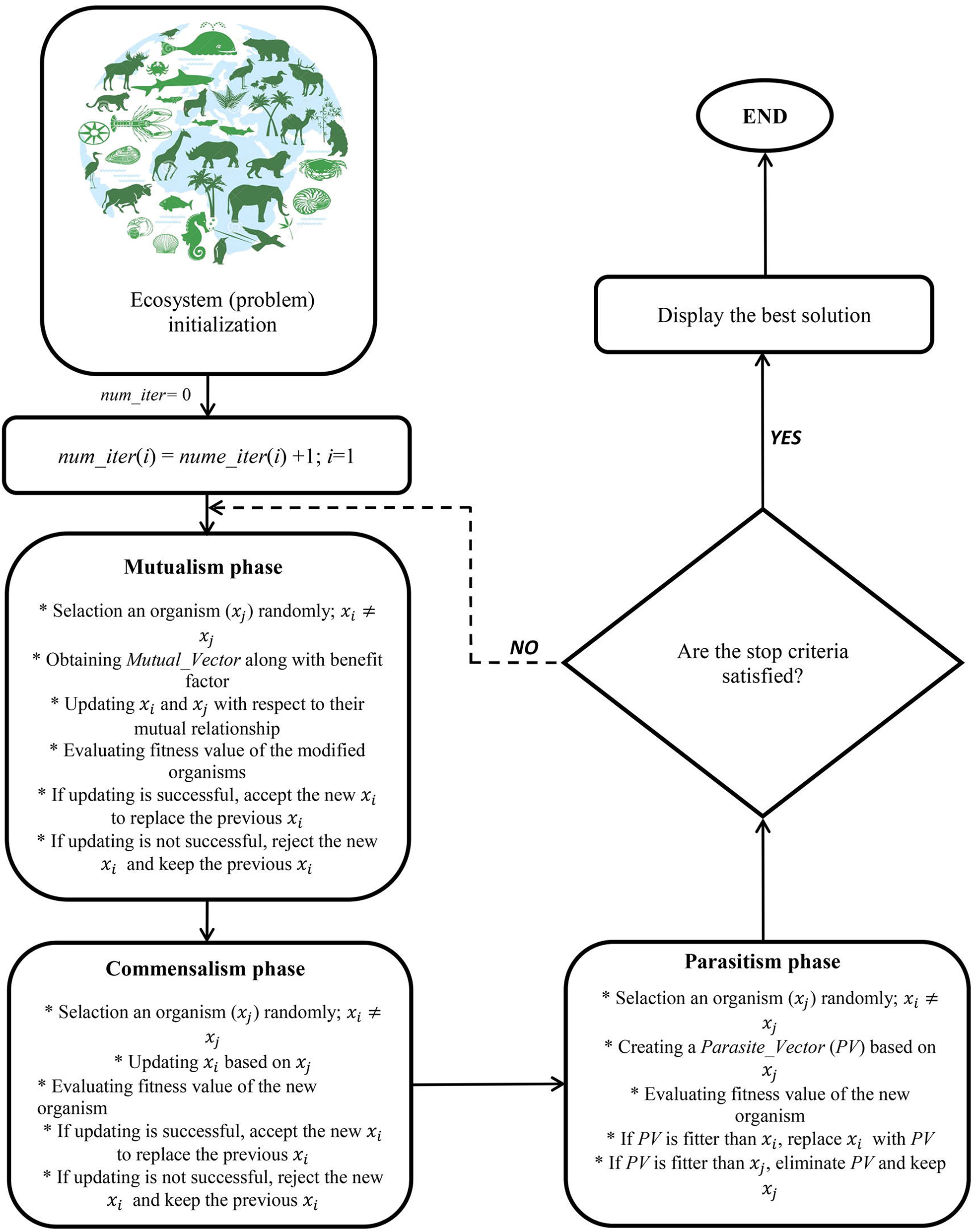
A schematic representation of symbiotic organisms search (SOS) algorithm.

After defining the maximal number of iterations and the number of species, the initial ecosystem is specified by generating a uniform random number between the upper and lower values of ecosystem size and a design variable (D) number. After that, *X*_*best*_ as the best current solution should be determined. In a process, named mutualism, two randomly chosen species along with *X*_*best*_ participate in a dialectic relationship that is profitable for both. New candidate solutions are generated based on the following equations:

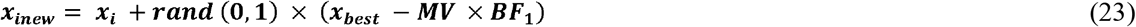

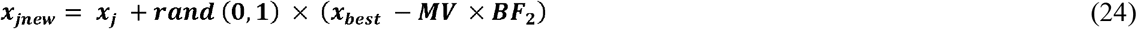

Where rand (0,1) shows a vector of random numbers, and the mutual vector (*MV*) equals the average value of *x*_*j*_ and *x*_*i*_ which enables the organisms to be updated concurrently rather than separately. In a mutualistic symbiosis between two species within nature, one species might gain a great advantage while the other receives no significant profit. This is presented by *BF*_*1*_ and *BF*_*2*_, which are randomly specified as either 1 or 2 ([*BF*_*i*_ = *Rand* (rand (0, 1) +1]; *i* = 1 and 2) to display the level of profits obtained from the relationship.

In the next step, the entire population is updated. Subsequently, the old candidate solutions *x*_*j*_ and *x*_*i*_ are compared with the new ones. More fit organisms are chosen as new solutions for the next iteration. The selections and comparisons start and end with the counter 1 and the counter equal to the *population size* (*npop*), respectively. For each *i*, the solution *j* is randomly chosen within the new population. Afterward, fitter organisms take part in the next step which is named commensalism. In commensalism, although one organism gains profits, the other remains neutral. Similar to the previous step, *x*_*j*_ is randomly chosen from the population to interact with *x*_*i*_. While *x*_*i*_ attempts to get profits from the engagement, *x*_*j*_ remains unaffected. If the new fitness value shows better performance than the previous one, the following equation is employed for updating xi:

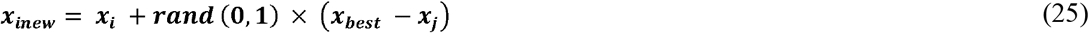

In the third step, which is named parasitism, the mutation operator of the SOS is required. In this step, *x*_*j*_ and *x*_*i*_ are the artificial host and parasite, respectively. In parasitism, one organism receives profits while the other is harmed. The sign of the parasite vector (*PV*) is that it competes with other randomly chosen dimensions instead of its parent with a series between upper and lower bounds. In this step, an initial parasite vector is produced by multiplying organism *x*_*j*_. Some of the decision variables from the parasite vector are randomly changed to recognize the parasite vector from *x*_*j*_. A random number should be produced in the range of [1, decision variable number] to describe the total number of changed variables. A uniform random number is produced for each dimension to achieve the position of the changed variables. Finally, a uniform distribution within the search area is needed for changing the variables and providing a parasite vector for the parasitism step. If the parasite vector displays better performance than *x*_*j*_ it becomes part of the population, whereas if *x*_*j*_ is not outperformed the parasite vector, *PV* eliminates from the population. The parasite vector is produced by changing *x*_*j*_ in random dimensions with random numbers rather than making small modifications in *x*_*j*_. If the current *x*_*j*_ and parasite vector are not the last member of the population, the SOS returns to the mutualism step that chosen *X*_*best*_ until obtaining a specified stopping criterion.

#### 2.4.4 Genetic algorithm (GA)

The GA, introduced by Holland (1992) is based on the Darwinian concepts of genetics and natural selection. Before applying the GA, some parameters such as crossover fraction, selection method, mutation rate, etc… should be specified. Subsequently, a set of possible answers are generated. The GA considers a set of chromosomes containing genes as an initial population. The genes represent the number of problem dimensions. During the optimization process, the genetic operators (e.g., Roulette Wheel and Tournament Selection) of the mutation and crossovers improve these genes.

Based on the competence of the chromosomes’ corresponding objective function, genes are selected to transfer to the next generation. The crossover operator replaces a number of genes from two chromosomes with each other. Moreover, the mutation operator changes some genes randomly. The elitism parameter is used to improve the chance of choosing the best chromosomes, then increase the convergence of the algorithm. When creating each new generation, three operators (i.e., crossover, selection, and mutation) regulate the optimization process in a way that the generated chromosomes improve the objective function value at each repetition until the optimization process will be completed by satisfying one of the termination criteria. The step-by-step procedure of the GA method has been presented in Figure 8.

**Figure 8.**
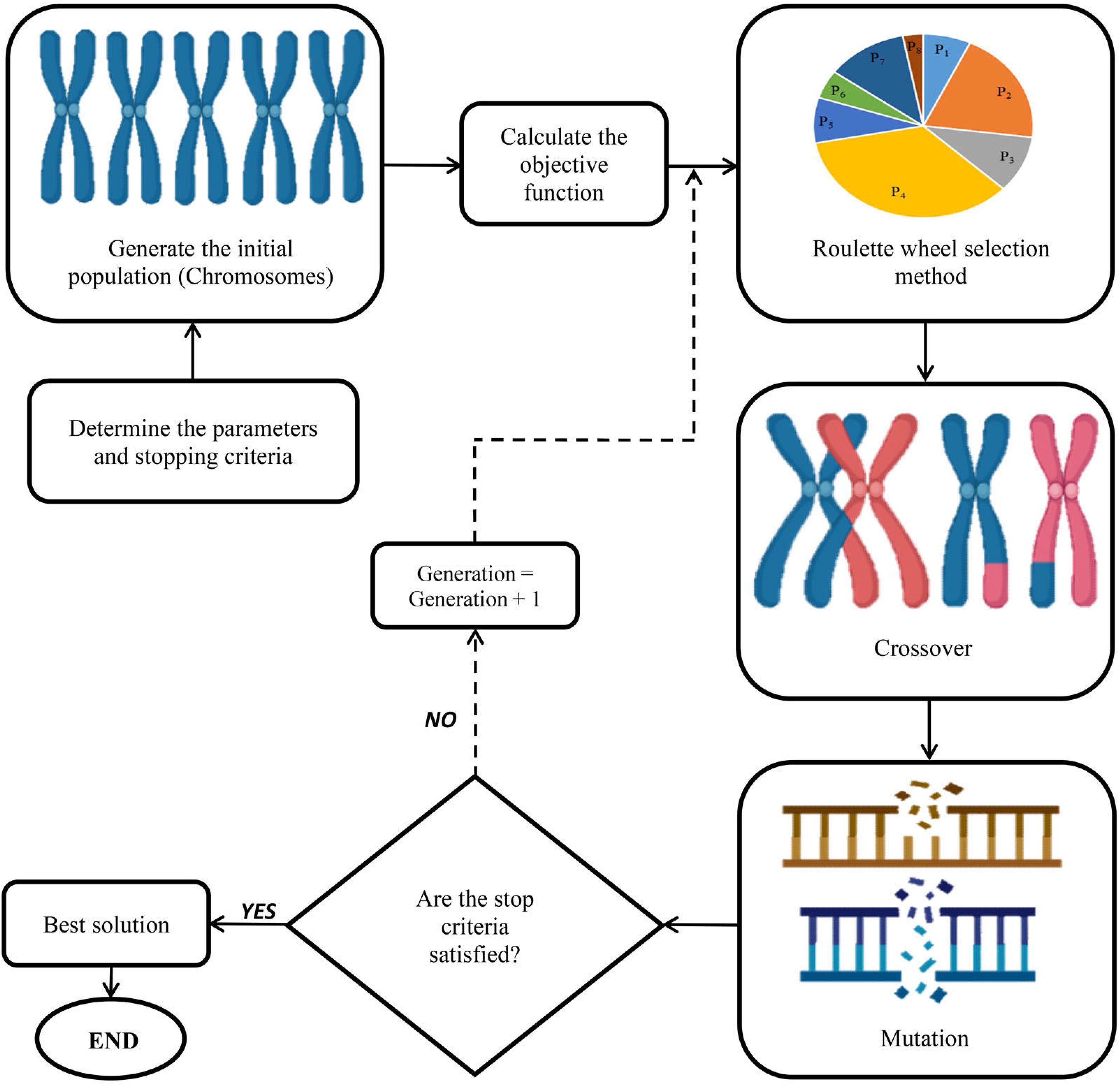
A schematic representation of genetic algorithm (GA).

### 1.5 Validation experiment

To evaluate the efficiency and reliability of the hybrid GRNN-evolutionary optimization algorithms, the predicted-optimized treatments obtained from evolutionary optimization algorithms (GA, ISA, SOS, and BBO) were separately evaluated in the lab as the validation experiment (Figure 9). The validation experiment was performed based on a completely randomized design with 4 replications. Effectiveness of optimized treatments were assessed by comparing error bars, representing standard error of means, as presented in Figure 4g,h.

**Figure 9.**
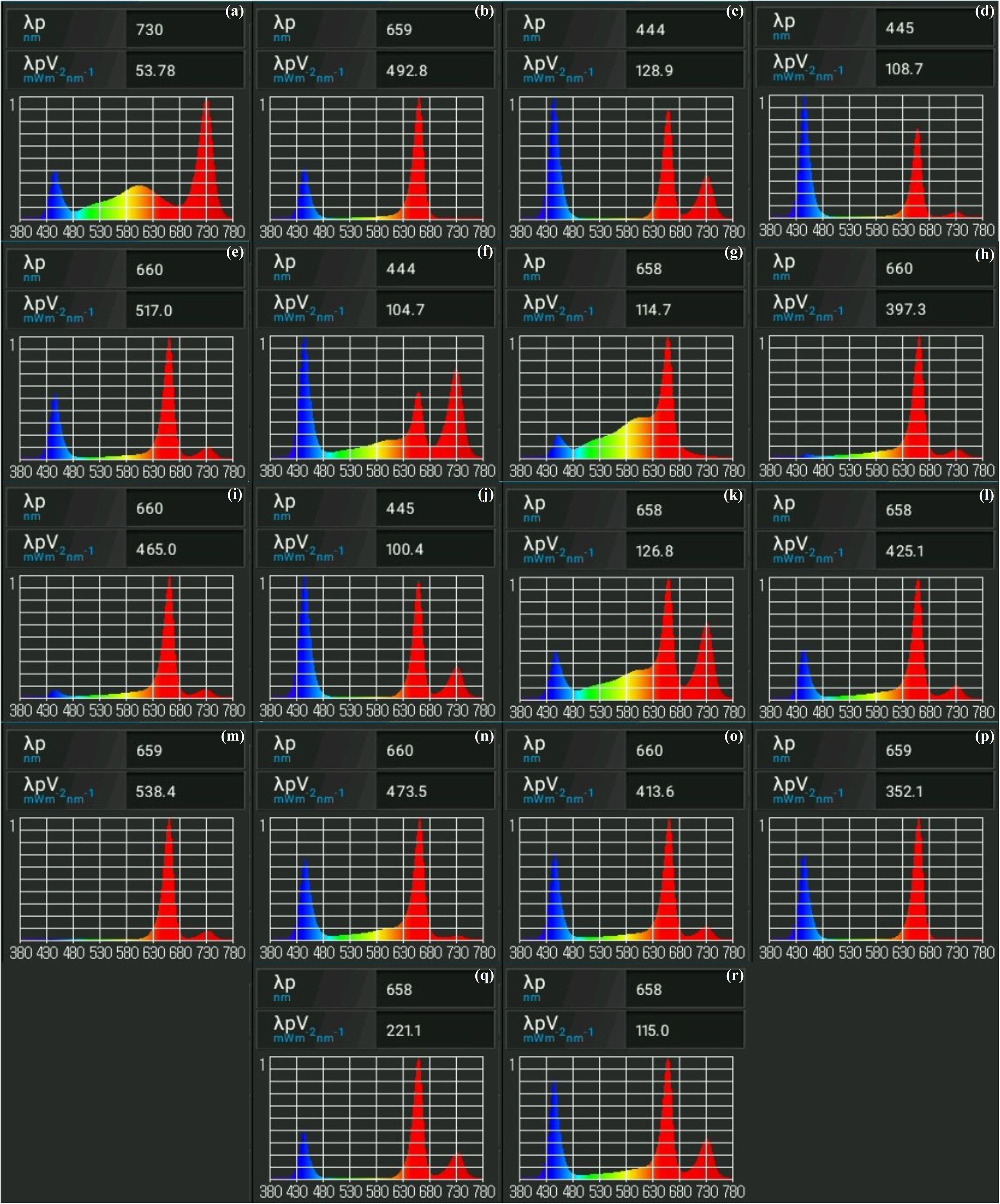
Spectral analyses of light treatments from the validation experiment. Images demonstrate relative amounts of fluencies emitted per treatment. Light spectra presented were obtained using Li-Cor LI-180 spectrometer. Presented are optimized light treatments for (a) GA shoot length, (b) GA canopy surface area, (c) GA number of shoots, (d) GA number of nodes, (e) BBO canopy surface area, (f) BBO shoot length, (g) ISA shoot length, (h) ISA number of nodes, (i) ISA root length, (j) ISA number of shoots, (k) SOS shoot length, (l) ISA canopy surface area, (m) SOS root length, (n) BBO number of nodes, (o) SOS canopy surface area, (p) SOS number of nodes, (q) BBO number of shoots, and (r) SOS number of shoots.

## 2 Results

### 2.1 Effects of light and carbohydrate sources on cannabis shoot growth and development

While this experiment was designed specifically for machine learning applications and standard statistical comparisons cannot be made, a wide range of responses were observed through the different treatments applied (Table 1). For instance, the greatest shoot length was acquired from 25 μmol/m^2^/s W + 25 μmol/m^2^/s Fr + 3 % Sucrose (154.68 ± 51.228 mm), while shoot length was most stunted when grown with 100 μmol/m^2^/s B + 1 % Sucrose (27.70 ± 2.311 mm). Greatest root length was achieved with 33.33 μmol/m^2^/s R + 33.33 μmol/m^2^/s B + 33.33 μmol/m^2^/s Fr + 3 % Sucrose (477.10 ± 287.094 mm), though no roots emerged from 100 μmol/m^2^/s B + 1 % Sucrose, or 50 μmol/m^2^/s W + 50 μmol/m^2^/s Fr + 1 % Sucrose specimens, and lowest root lengths were observed from those of the 100 μmol/m^2^/s W + 1 % Sucrose (4.36 ± 4.363 mm) treatment. Plantlets developing the most nodes came from 33.33 μmol/m^2^/s R + 33.33 μmol/m^2^/s B + 33.33 μmol/m^2^/s Fr + 3 % Sucrose (11.50 ± 2.901), while the fewest nodes were observed in 100 μmol/m^2^/s B + 1 % Sucrose (5.75 ± 0.479) treated plantlets. The largest canopy surface area was attained by plantlets grown under 50 μmol/m^2^/s R + 50 μmol/m^2^/s B + 3 % Sucrose (13061.97 ± 10839.642 mm^2^), whereas smallest canopy was observed in 100 μmol/m^2^/s B + 1 % Sucrose (493.01 ± 111.615 mm^2^), 100 μmol/m^2^/s W + 1 % Sucrose (519.06 ± 182.411 mm). Results for the preliminary experiment are outlined in Table 1. Treatments consisted of single experimental units with 4 biological replicates each, which satisfied the models used, with high accuracy.

Based on our observations (Table 1), a general trend is observed for appreciable shoot length when sucrose concentration is 3% (w/v), irradiance levels are in range of 50-100 μmol/m^2^/s, and when W is included in multi-spectral treatments. These treatments allowed long shoot length that developed between 32.44 ± 7.036 - 154.68 ± 51.228mm. Additionally, there was a broad tendency for multi-spectral treatments with 75-100 μmol/m^2^/s that included R and 3% (w/v) sucrose to develop large canopy surface areas, which ranged from 2483.71 ± 627.011 - 13061.97 ± 10839.642 mm^2^ (Table 1).

Of the 66 treatments tested, 50 μmol/m^2^/s W + 50 μmol/m^2^/s Fr + 1 % Sucrose noticeably accumulated phenolic compounds in the media, which was not observed in any other treatment. Additionally, 12 cultures produced plantlets with floral organs despite being grown under a long day photoperiod. Cultures included 25 μmol/m^2^/s B + 25 μmol/m^2^/s W + 1 % Sucrose, 25 μmol/m^2^/s R + 75 μmol/m^2^/s B + 1 % Sucrose, 25 μmol/m^2^/s R + 75 μmol/m^2^/s B + 3 % Sucrose, 33.33 μmol/m^2^/s R + 33.33 **+**33.33 Fr + 1 % Sucrose, 33.33 R + 33.33 B + 33.33 Fr + 3 % Sucrose, 25 μmol/m^2^/s R + 75 μmol/m^2^/s B + 6 % Sucrose, 25 μmol/m^2^/s R + 25 μmol/m^2^/s B + 25 μmol/m^2^/s Fr + 25 μmol/m^2^/s W + 6 % Sucrose, 25 μmol/m^2^/s R + 25 μmol/m^2^/s B + 6 % Sucrose, 100 μmol/m^2^/s B + 6 % Sucrose, 50 μmol/m^2^/s B + 6 % Sucrose, 25 μmol/m^2^/s B + 25 μmol/m^2^/s W + 6 % Sucrose, 50 μmol/m^2^/s + 6 % Sucrose, 50 μmol/m^2^/s R + 50 μmol/m^2^/s B + 3 % Sucrose, 50 μmol/m^2^/s W + 50 μmol/m^2^/s Fr + 3 % Sucrose.

50 μmol/m^2^/s B treatments, for the most part, showed higher values relating to developmental features than 100 μmol/m^2^/s B (Table 1). This is likely due to malfunctioning 100 μmol/m^2^/s B lights, which were repaired within a few days. However, we nonetheless attribute the delayed development of 100 μmol/m^2^/s B treatments to the brief period of light malfunction.

### 2.2 Data modeling through MLP, GRNN, and ANFIS

Machine learning algorithms including MLP, GRNN, and ANFIS were employed to model and predict cannabis shoot growth and development traits (shoot length, root length, number of nodes, number of shoots, and canopy surface area) as target variables based on five input variables (Sucrose, B, R, W, and Fr). R^2^, RMSE, and MBE were used to assess the prediction performance of the developed machine learning algorithms (Table 2). The GRNN model presented higher R^2^ as one of the most important performance indices in comparison to MLP or ANFIS in both training and testing processes for all shoot growth and development traits including shoot length (*R*^*2*^ > 0.96 for GRNN vs. *R*^*2*^ > 0.58 for ANFIS or *R*^*2*^ > 0.95 for MLP), root length (*R*^*2*^ > 0.91 for GRNN vs. *R*^*2*^ > 0.58 for ANFIS or *R*^*2*^ > 0.89 for MLP), number of nodes (*R*^*2*^ > 0.74 for GRNN vs. *R*^*2*^ > 0.54 for ANFIS or *R*^*2*^ > 0.39 for MLP), number of shoots (*R*^*2*^ > 0.71 for GRNN vs. *R*^*2*^ > 0.50 for ANFIS or *R*^*2*^ > 0.42 for MLP), and canopy surface area (*R*^*2*^ > 0.94 for GRNN vs. *R*^*2*^ > 0.64 for ANFIS or *R*^*2*^ > 0.92 for MLP) (Table 2). Also, higher RMSE and MBE for GRNN in comparison to MLP and ANFIS for all studied traits indicated that the assessed results were highly accurate and correlated, showing the good performance of the developed GRNN models (Table 2). Moreover, the regression lines displayed a good fit correlation between experimental and predicted data for all the shoot growth and development traits in both training and testing processes (Fig. 10).

**Table 2.** Performance indices of different machine learning algorithms (MLP, GRNN, and ANFIS) for modeling and predicting shoot length, root length, number of nodes, number of shoots, and canopy surface area of *Cannabis*.

| Model | Performance index | Shoot length |  | Shoot number |  | Node number |  | Root length |  | Canopy surface area |  |
| --- | --- | --- | --- | --- | --- | --- | --- | --- | --- | --- | --- |
|  |  | Training | Testing | Training | Testing | Training | Testing | Training | Testing | Training | Testing |
| MLP | R<sup>2</sup> | 0.972 | 0.954 | 0.625 | 0.421 | 0.717 | 0.390 | 0.938 | 0.900 | 0.953 | 0.928 |
|  | RMSE | 4.929 | 6.927 | 0.396 | 0.632 | 0.702 | 1.202 | 15.112 | 15.956 | 277.487 | 340.538 |
|  | MBE | -0.090 | 1.673 | 0.009 | -0.001 | 0.017 | 0.260 | 0.001 | 2.259 | 30.952 | 25.016 |
| GRNN | R<sup>2</sup> | 0.983 | 0.964 | 0.733 | 0.714 | 0.791 | 0.744 | 0.941 | 0.914 | 0.962 | 0.944 |
|  | RMSE | 3.879 | 6.081 | 0.347 | 0.606 | 0.594 | 0.933 | 14.754 | 14.972 | 248.737 | 300.911 |
|  | MBE | 0.001 | 1.540 | 0.001 | 0.012 | -0.001 | 0.063 | 0.001 | 2.581 | 0.001 | 2.388 |
| ANFIS | R<sup>2</sup> | 0.770 | 0.590 | 0.647 | 0.501 | 0.767 | 0.549 | 0.781 | 0.589 | 0.733 | 0.644 |
|  | RMSE | 17.538 | 23.327 | 0.407 | 0.557 | 0.650 | 0.942 | 41.881 | 39.007 | 1282.011 | 1282.697 |
|  | MBE | -4.549 | -5.508 | 0.006 | -0.065 | -0.003 | 0.037 | 5.962 | 8.546 | -26.525 | -32.037 |

ANFIS: adaptive neuro-fuzzy inference system; GRNN: generalized regression neural network; MBE: mean bias error; MLP: multi-layer perceptron; R<sup>2</sup>: coefficient of determination; RMSE: root mean square error.

**Figure 10.**
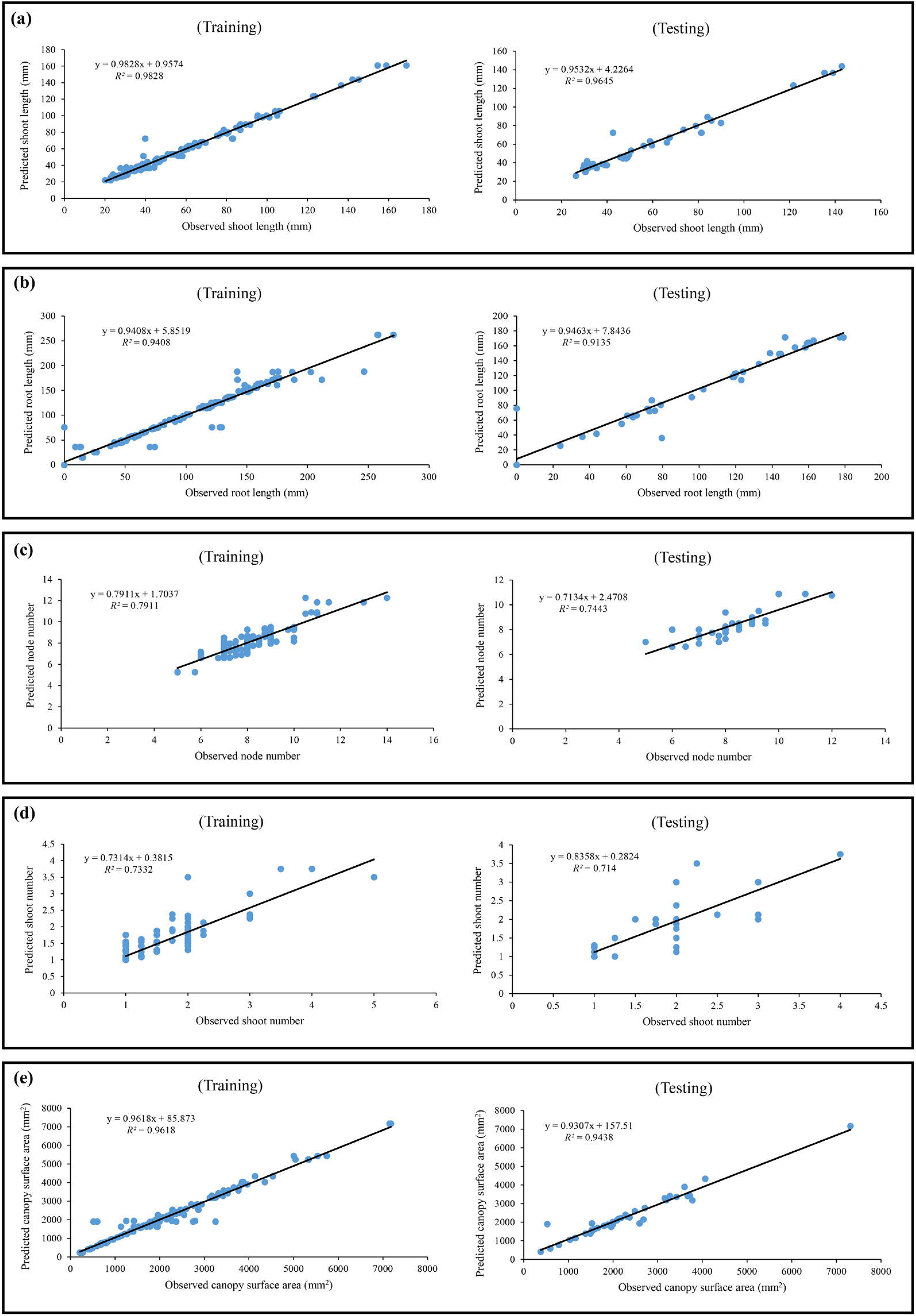
Scatter plot of experimental data versus predicted data of (a) shoot length, (b) root length, (c) node number, (d) shoot number, and (e) canopy surface area in *in vitro Cannabis* shoot growth and development, using generalized regression neural network (GRNN) in both training and testing subsets.

### 2.3 Determining the importance of each input on cannabis shoot growth and development

To determine the importance of each input variable on the objective function (studied parameter including shoot length, root length, number of nodes, number of shoots, and canopy surface area) sensitivity analysis was performed by calculating VSR. The results showed that both shoot length and node number were more sensitive to Sucrose followed by B, Fr, R, and W, while root length was more sensitive to Sucrose followed by R, Fr, W, and B light (Table 3). Also, the results demonstrated more sensitivity of shoot number to Sucrose followed by B, red, Fr, and W color (Table 3). Moreover, Sucrose> R> B> W> Fr were ranked for canopy surface area (Table 3).

**Table 3.** Importance degree of light (blue, red, white, and far-red) and carbohydrate sources on shoot length, root length, number of nodes, number of shoots, and canopy surface area of *Cannabis* through sensitivity analysis.

| Trait | Item | Blue | Red | White | Far-red | Sucrose |
| --- | --- | --- | --- | --- | --- | --- |
| Shoot length | VSR | 3.005 | 1.957 | 1.647 | 2.141 | 5.191 |
|  | Rank | 2 | 4 | 5 | 3 | 1 |
| Root length | VSR | 1.54 | 2.211 | 1.669 | 1.909 | 3.887 |
|  | Rank | 5 | 2 | 4 | 3 | 1 |
| Node number | VSR | 1.379 | 1.257 | 1.065 | 1.288 | 1.597 |
|  | Rank | 2 | 4 | 5 | 3 | 1 |
| Shoot number | VSR | 1.554 | 1.217 | 1.105 | 1.168 | 1.651 |
|  | Rank | 2 | 3 | 5 | 4 | 1 |
| Canopy surface area | VSR | 1.662 | 2.616 | 1.657 | 1.622 | 3.693 |
|  | Rank | 3 | 2 | 4 | 5 | 1 |

VSR: variable sensitivity ratio

### 2.4 Optimization process via GA, SOS, ISA, and BBO

In the present study, four different evolutionary optimization algorithms including BBO, ISA, SOS, and GA were separately used to determine the optimal level of Sucrose, B, R, W, and Fr for maximizing each fitness function (shoot length, root length, number of nodes, number of shoots, and canopy surface area). Although all optimization algorithms predicted the same best fitness function value, they found a different optimal level of inputs for each fitness function (Table 4). For instance, the maximum shoot length (160.78 mm) would be achieved from 15.412 μmol/m^2^/s B + 9.412 μmol/m^2^/s R + 15.997 μmol/m^2^/s W + 43.271 μmol/m^2^/s Fr + 3.142 % Sucrose based on the BBO, 4.460 μmol/m^2^/s B + 19.051 μmol/m^2^/s R + 27.337 μmol/m^2^/s W + 39.472 μmol/m^2^/s Fr light + 3.157 % Sucrose based on the SOS, 0.439 μmol/m^2^/s B + 18.494 μmol/m^2^/s R + 36.234 μmol/m^2^/s W + 33.122 μmol/m^2^/s Fr + 3.319 % Sucrose based on the ISA, or 2.937 μmol/m^2^/s B + 0.270 μmol/m^2^/s R + 13.036 μmol/m^2^/s W + 30.605 μmol/m^2^/s Fr light + 3.505 % Sucrose based on the GA (Table 4). Also, 52.563 μmol/m^2^/s B + 84.052 μmol/m^2^/s R + 26.262 μmol/m^2^/s W + 22.456 μmol/m^2^/s Fr light + 3.809 % Sucrose based on the BBO, 54.688 μmol/m^2^/s B + 95.974 μmol/m^2^/s R + 30.099 μmol/m^2^/s W + 24.543 μmol/m^2^/s Fr light + 3.664 % Sucrose based on the SOS, 44.889 μmol/m^2^/s B + 99.642 μmol/m^2^/s R + 49.994 μmol/m^2^/s W + 24.674 μmol/m^2^/s Fr light + 3.285% Sucrose based on the ISA, or 37.646 μmol/m^2^/s B + 83.928 μmol/m^2^/s R + 17.507 μmol/m^2^/s W + 1.811 μmol/m^2^/s Fr + 3.083 % Sucrose based on the GA would result in the highest canopy surface area (7168.05 mm^2^) (Table 4).

**Table 4.** The results of optimization process via different evolutionary optimization algorithms (BBO, SOS, ISA, and GA).

| Fitness function | Optimization algorithm | Optimal level of input variables |  |  |  |  | Predicted fitness function value |
| --- | --- | --- | --- | --- | --- | --- | --- |
| | | Blue<br>($\mu\text{mol/m}^2/\text{s}$) | Red<br>($\mu\text{mol/m}^2/\text{s}$) | White<br>($\mu\text{mol/m}^2/\text{s}$) | Far-red<br>($\mu\text{mol/m}^2/\text{s}$) | Sucrose<br>(%) | |
| Shoot length (mm) | BBO | 15.412 | 9.412 | 15.997 | 43.271 | 3.142 | 160.78 |
|  | SOS | 4.460 | 19.051 | 27.337 | 39.472 | 3.157 | 160.78 |
|  | ISA | 0.439 | 18.494 | 36.234 | 33.122 | 3.319 | 160.78 |
|  | GA | 2.937 | 0.270 | 13.036 | 30.605 | 3.505 | 160.78 |
| Root length (mm) | BBO | 5.756 | 87.381 | 31.523 | 19.343 | 3.504 | 262.21 |
|  | SOS | 0.508 | 79.897 | 4.733 | 17.209 | 3.673 | 262.21 |
|  | ISA | 3.779 | 81.519 | 25.386 | 13.321 | 3.634 | 262.21 |
|  | GA | 5.797 | 98.198 | 36.208 | 1.237 | 3.507 | 262.21 |
| Node number | BBO | 62.998 | 92.238 | 48.520 | 8.830 | 3.709 | 12.25 |
|  | SOS | 57.682 | 87.192 | 45.468 | 19.924 | 3.711 | 12.25 |
|  | ISA | 46.845 | 90.135 | 20.969 | 16.824 | 3.220 | 12.25 |
|  | GA | 50.960 | 88.355 | 11.000 | 11.673 | 3.709 | 12.25 |
| Shoot number | BBO | 16.581 | 36.686 | 0.592 | 19.995 | 2.909 | 3.75 |
|  | SOS | 15.930 | 21.183 | 9.723 | 24.304 | 2.372 | 3.75 |
|  | ISA | 21.336 | 29.584 | 0.332 | 17.387 | 2.174 | 3.75 |
|  | GA | 25.303 | 25.471 | 0.262 | 18.125 | 3.160 | 3.75 |
| Canopy surface area<br>($\text{mm}^2$) | BBO | 52.563 | 84.052 | 26.262 | 22.456 | 3.809 | 7168.05 |
|  | SOS | 54.688 | 95.974 | 30.099 | 24.543 | 3.664 | 7168.05 |
|  | ISA | 44.889 | 99.642 | 49.994 | 24.674 | 3.285 | 7168.05 |
|  | GA | 37.646 | 83.928 | 17.507 | 1.811 | 3.083 | 7168.05 |

BBO: biogeography-based optimization; GA: genetic algorithm; ISA: interior search algorithm; SOS: symbiotic organisms search.

### 2.5 Determining the reliability of the developed models

The optimized-predicted results from each evolutionary optimization algorithm for shoot length and canopy surface area as fitness functions were experimentally tested in a validation experiment to evaluate the reliability of the developed models. Based on the validation experiment results, the differences among evolutionary optimization algorithms (GA, ISA, BBO, and SOS) and optimized-predicted results for both shoot length (Fig. 4f) and canopy surface area (Fig. 4g) were negligible, which demonstrated the reliability of the developed models. However, the maximum shoot length (206.76 ± 41.542 mm) (Fig. 4h) and canopy surface area (8193.49±2102.624 mm^2^) (Fig. 4i) were achieved from the GRNN-SOS, while GRNN-BBO resulted in the lowest shoot length (181.83 ± 39.676 mm) and canopy surface area (5745.34 ± 919.848 mm^2^). Therefore, it seems that the SOS has better performance than the other optimization algorithms.

## 3 Discussion

As with any *in vitro* culture system, many intrinsic (e.g., genotype, type and age of explant) and extrinsic (e.g., basal salt medium, vitamins, PGRs, gelling agent, carbohydrate source, additives, temperature, and light) factors influence *in vitro* shoot growth and development. Fortunately, due to the highly controlled nature of plant tissue culture, most of these factors can be manipulated to evaluate their impact on system optimization. Historically, micropropagation systems were refined using traditional statistical models to sequentially manipulate and optimize single factors. This approach often requires hundreds or even thousands of treatments to be tested, and even then sequential optimization does not account for interactions and can miss the best combinations (García-Pérez et al., 2020; Hameg et al., 2020). Due to the cost and time requirements, most species are cultured in conditions optimized for other species with minor modifications and are not fully optimized for any given application. In our study, we demonstrate that specific growth responses of *in vitro* cannabis can be directed by manipulating abiotic factors such as light intensity, spectrum, and exogenous carbon availability, and that machine learning approaches provide an effective approach to optimize these factors for specific outcomes. It is possible that these modifications could trigger developmental changes by regulating photosynthetic activity (Hdider and Desjardins, 1994), or by regulating intrinsic concentrations of phytohormones (Premkumar et al., 2001). Additional experiments must be completed to indicate the precise mechanisms by which dynamic physiological responses occur. Ultimately, we clearly show that plant growth and development can be influenced by light quality and sucrose levels in the absence of PGRs.

Efficient protocol development is a long-standing challenge in the field and more advanced statistical models using surface response curves have been applied with some success (Niedz and Evens, 2016; Niedz and Marutani-Hert, 2018; Pence et al., 2020). While these methods are more efficient than sequential optimization and can account for interactions among factors, they are still limited in the number of factors that can be included in a single experiment, require several assumptions to be met that are often not possible to achieve, require significant numbers of treatments, and rely on relatively simple interactions that can be compared using regression analyses. An alternative to address the inherent complexity of plant tissue culture systems is to apply machine learning methodology. This approach leverages modern computing power and developments in artificial intelligence to efficiently recognize patterns in complex and disorderly datasets, typical of what is observed in plant tissue culture (Hesami et al., 2021; Hesami and Jones, 2020). Machine learning algorithms can then be combined with optimization algorithms to decipher complex interactions and predict theoretically optimized combinations of factors for desired outcomes. The combination of machine learning and optimization algorithms has the potential to overcome many of the challenges associated with optimizing *in vitro* plant systems and enable development of more effective protocols using fewer treatments. Ultimately, this approach can be used to change the face of plant tissue culture advancements by enhancing the viability of optimization for specific species, or even individual genotypes.

Here, ANNs (MLP and GRNN) and neuro-fuzzy logic (ANFIS) were employed and compared to model and predict the effects of light quality and carbohydrate supply on growth and development of *in vitro* cannabis plants. Based on our results, using the stated parameters, GRNN had better performance than either MLP or ANFIS. Although there are no studies in plant tissue culture comparing the predictive performances of neuro-fuzzy logic systems and ANNs, several studies in other fields have demonstrated that GRNN often performs better than MLP or ANFIS. For instance, Sridharan (2021) reported that the prediction accuracy of GRNN was better than MLP and ANFIS for modeling and predicting global solar irradiance. Similar results were also reported by Ausati and Amanollahi (2016) who showed GRNN performed better than ANFIS and MLP for modeling and predicting air pollution.

In the present study, four evolutionary optimization algorithms (BBO, GA, ISA, and SOS) were individually linked to the GRNN to determine optimal levels of Sucrose, B, R, W, and Fr for maximizing each fitness function (shoot length, root length, number of nodes, number of shoots, and canopy surface area). Based on mean standard errors reported in our results, there is no difference in the predicted values of fitness functions among different optimization algorithms. Although the results of the validation experiments showed that the differences in the performance of the optimization algorithms were negligible, SOS led to the highest level of studied fitness functions. For instance, GRNN-SOS showed that using the theoretical optimal combination of light quality and sucrose levels, average shoot length and canopy surface area were 206.76 ± 41.542 mm and 8193.49 ± 2102.624 mm^2^, respectively. Although no studies exist for using and comparing dissimilar optimization algorithms for *in vitro* culture optimization, several studies previously showed that SOS can be considered one of the most powerful of the evolutionary optimization approaches (Bozorg-Haddad et al., 2016; Cheng and Prayogo, 2014). Bozorg-Haddad et al., (2016) compared GA and SOS for optimization of reservoir operation. They run these two algorithms 10 times and reported that there was no significant difference between the performances of GA and SOS, however, SOS calculated higher fitness function values than GA in all 10 runs. Similar to this result, the results of our validation experiment showed that SOS resulted in a higher value of fitness function in comparison with other algorithms.

Based on the sensitivity analysis, sucrose was the most important factor for all traits studied (shoot length, root length, shoot number, node number, and canopy surface area). This likely reflects the mixotrophic nature of *in vitro* plants and limitations the sealed environment (depletion of CO2, high relative humidity, etc.) places upon their photosynthetic capacity (De La Viña et al., 1999; Nguyen et al., 2001; Shin et al., 2013). Due to these limitations, supplemental sucrose appears to be critical to support plant growth and development. It is likely that different results may be obtained if this experiment were conducted using vented lids or forced air, which would improve potential photosynthesis and increase the relative importance of light quality.

In our experiment, evolutionary optimization algorithms predicted that ~3 % sucrose would result in the highest shoot growth and development. A plethora of previous studies have found 2–4 % sucrose, in particular 3 % (w/v), to be optimal for various species and this has become a standard for most micropropagation systems (reviewed by Yaseen et al., 2013). For instance, the results of GRNN-SOS showed that 3.157 % sucrose would lead to the highest shoot length. Similar to our results, Romano et al. (1995) and Baskaran and Jayabalan (2016) respectively studied different levels of *in vitro* sucrose on shoot growth and development of *Quercus suber* L. and *Eclipta alba* (L.) Hassk. They reported that 3% sucrose was optimal for maximizing shoot length *in vitro*. Although the effect of sucrose concentration on cannabis micropropagation needs more attention, previous reports generally use 3% (w/v) for shoot growth and development (reviewed by Hesami et al., 2021). These results support the standard use of 3% sucrose for micropropagation, but more importantly demonstrate the ability of machine learning techniques to optimize environmental factors in tissue culture systems using a relatively small number of treatments.

Though sucrose is identified as the most important factor in plant growth and development in this study, light intensity and spectrum also play important roles for *in vitro* morphogenic and developmental processes (Batista et al., 2018). Different photoreceptors recognize the quality and quantity of light (e.g., phytochromes absorb red and far-red, phtotropins and cryptochromes absorb blue light), and subsequently use this information to direct photomorphogenic functions (Li et al., 2012; Parihar et al., 2016). Several studies have previously shown the impact of light quality and quantity on different tissue culture systems for shoot growth and development (Hung et al., 2016), somatic embryogenesis (Ferreira et al., 2017; Hesami et al., 2019), rhizogenesis (Gago et al., 2014), and secondary metabolite production (Dutta Gupta and Karmakar, 2017; Silva et al., 2017). However, each *in vitro* developmental stage requires a specific light condition (Batista et al., 2018). Our sensitivity analysis showed that, among light treatments, B was the most important factor for shoot length, shoot number, and node number, while R had the highest degree of importance on root length and canopy surface area. The importance of B and R on *in vitro* shoot growth and development has been previously confirmed in different plants such as *Myrtus communis* L. (Cioć et al., 2018), *Plectranthus amboinicus* (Lour.) Spreng (Silva et al., 2017), *Pfaffia glomerata* (Spreng.) Pedersen (Silva et al., 2020), *Achillea millefolium* L. (Alvarenga et al., 2015), and *Stevia rebaudiana* Bertoni (Ramírez-Mosqueda et al., 2017).

Light intensity is another important parameter that should be optimized for each *in vitro* culture stage. Through GRNN-SOS, the predicted optimal spectrum included 4.460 μmol/m^2^/s B + 19.051 μmol/m^2^/s R + 27.337 μmol/m^2^/s W + 39.472 μmol/m^2^/s Fr light + 3.157 % Sucrose to maximize shoot length. In total, this provides about 50.8 μmol/m^2^/s PAR plus 39.472 μmol/m^2^/s Fr. In line with our results, Silva et al. (2017) reported that light intensity below 51 μmol/m^2^/s resulted in the highest shoot length in *P*. *amboinicus*. Similar results were also reported by Alvarenga et al (2015) for *A*. *millefolium*. However, using GRNN-SOS to predict the optimal spectrum to maximize canopy surface area, the conditions included 54.688 μmol/m^2^/s B + 95.974 μmol/m^2^/s R + 30.099 μmol/m^2^/s W + 24.543 μmol/m^2^/s Fr + 3.664 % sucrose, for a total of 180.8 μmol/m^2^/s PAR plus 24.5 μmol/m^2^/s Fr. Alternatively, GRNN-BBO conditions included 52.563 μmol/m^2^/s B + 84.052 μmol/m^2^/s R + 26.262 μmol/m^2^/s W + 22.465 μmol/m^2^/s Fr + 3.809 % sucrose, totaling 162.9 PAR + 22.5 μmol/m^2^/s Fr, which ultimately resulted in the lowest canopy surface areas. Here, the difference between GRNN-SOS and GRNN-BBO relating to Fr fluence is 2.1 μmol/m^2^/s, while the total difference in PAR fluence is 17.9 μmol/m^2^/s, 11.9 μmol/m^2^/s of which is the dissimilarity of R intensity. This leads us to speculate that PAR intensity, specifically R, is an important factor governing canopy development. This is confirmed with the sensitivity analysis ranking, as R is the most important spectra for this growth parameter. Alternative wavelengths of light can be efficiently absorbed at different depths within the leaf tissue. This can also be enhanced with increasing light intensity. While certain wavelengths of green light can penetrate deeper into leaves, R and B can effectively be absorbed toward the leaf surface (Zheng and Van Labeke, 2017), triggering phytochrome and cryptochrome –mediated re-localization of phytohormones for photo-morphogenesis (Miler and Zalewska, 2006). Although the optimal light intensity varies by species, most micropropagation systems use light levels ranging from 40-80 μmol/m^2^/s PAR (Miler et al., 2019; Murphy and Adelberg, 2021; Nhut et al., 2003). However, some species perform better at higher fluence rates, for example, *Actinidia deliciosa* (Gago et al., 2014), *Lippia gracilis* (Lazzarini et al., 2018), and *Solanum tuberosum* (Kulchin et al., 2018). In general, cannabis is known to grow best *in vivo* under higher light levels (Murphy and Adelberg, 2021; Wróbel et al., 2020), with yields increasing linearly up to at least 1600 μmol/m^2^/s PAR (Chandra et al., 2008; Lata et al., 2016; Rodriguez-Morrison et al., 2021), depending on the culture system. As such, the prediction to use such high light levels may reflect the nature of the species. Our validation experiment demonstrates that cannabis responded to these high light levels as predicted.

Similarly, in our initial experiment, we observed higher intensities of R in combination with equal or lower intensity of B or W to be beneficial to canopy development in the presence of 3 % Sucrose. Unlike some other treatments, 50 μmol/m^2^/s R + 50 μmol/m^2^/s B + 3 % Sucrose, with the largest canopy surface area, did not give largest averages in any additional growth parameters measured. This is counterintuitive on the premise that canopy surface area is a metric of leaf size in addition to leaf number. We might expect highest canopy surface area treatments to be mutually high in other shoot growth parameters such as shoot length, number of nodes, or number of shoots. R significantly impacts endogenous action of gibberellic acid which is involved in cell elongation, root inhibition, and stimulating mitosis in meristematic cells (Manivannan et al., 2015) for replication. Gibberellic acid action is known to trigger anisotropic responses for leaf expansion in monocots (Sprangers et al., 2020; Xu et al., 2016), though R can impart different effects on leaf morphology for different plants *in vitro*. B increased leaf thickness, leaf numbers and leaf areas compared to R, which reduced leaf thickness and area in cultured *Alternanthera brasiliana* (Macedo et al., 2011). Similarly, B mutually amplified leaf thickness and leaf area of *Ficus benjamina* (Zheng and Van Labeke, 2017), and *Cucumis sativus in vivo*, as well as micropropagated *Solanum tuberosum* L. (Chen et al., 2020). B also had a tendency to increase leaf area of *Cordyline australis* and *Sinningia speciosa in vivo* (Zheng and Van Labeke, 2017). Since we observed opposite influences of B, we can speculate that influences of this spectrum to be species-dependent. Wei et al. (2021) found that LED-treated hemp plants produced smaller leaf areas than high pressure sodium treatments, though the LED treatments with higher R:B at higher intensities produced larger leaf areas than treatments of lower R:B ratios at higher or lower intensities. They also found leaf areas to bear a significantly positive correlation leaf number, though no additional growth responses or treatments were significantly correlated with leaf area (Wei et al., 2021). These results correspond more similarly with the data obtained in our study, though it’s difficult to imply for certain if *in vitro* medicinal cannabis responds to light quality and intensity with the same general trend when influenced by sucrose in a sub-optimal gaseous environment. It is also difficult to infer molecular mechanisms for such *in vitro* plant responses, since they are beyond the scope of the presented study. Thus, subsequent experiments should be devised to test molecular mechanisms of the growth parameters measured to further elucidate the molecular devises contributing to the factors observed.

Our preliminary experiment also indicated shade avoidance-like responses observed when comparing R + B + Fr + 3 % Sucrose treatments at different light intensities. Higher light intensity generally produced shorter shoots with more nodes versus longer shoots with fewer nodes when irradiance was lower. At higher light intensities, R + B + Fr + 6 % Sucrose specimens also averaged longer stems with more nodes than at 1% Sucrose, though at low light intensity R + B + Fr + 1 % Sucrose grew longer stems with more nodes than 6 % plants of the same light treatment. These observations imply that that there is a complex interaction between sugar and light signaling whereby the impact of sugar can allow plants to dynamically adjust to higher light intensities (Tichá et al., 1998), or impede certain physiological responses when exogenous carbon is too high and abiotic factors are sub-optimal (Roh and Choi, 2004). However, in all cases, greatest averages were achieved with 3 % sucrose, which suggests that sugar concentrations above 4 % and below 2 % can have diminishing returns on shoot development and number (Sivanesan and Park, 2015). This observation supports the widespread practice of using 3% sucrose in plant tissue culture systems, and the results of our sensitivity analysis. Gago et al. (2014) modeled 14 growth parameters of *in vitro* kiwifruit based on Sucrose concentration and irradiance, using Neuro-fuzzy logic. They found an *in vitro* sucrose concentration of 2.3 % or higher to be indispensable in achieving many of the optimal growth parameters investigated, either independently or in interaction with light intensity. Dynamic interactions between light and exogenous sugar are important for evoking many additional physiological responses relating to light attenuation and metabolism (Gago et al., 2014; Roh and Choi, 2004; Tichá et al., 1998).

Lalge et al. (2017) observed that taller cannabis clones developed with W compared to B + R LEDs when grown in controlled climates. The optimal levels of R:Fr in promoting stem elongation has been well documented (Ballaré and Pierik, 2017; Ma and Li, 2019; Trupkin et al., 2014). Though B also impacts stem growth (Ma and Li, 2019; Magagnini et al., 2018; Snowden et al., 2016), it can sometimes have the opposite influence of R:Fr, resulting in more compact phenotypes (Magagnini et al., 2018). The optimized combinations R:Fr in addition to B could have ultimately impacted shoot elongation of the treatments assessed (Cope and Bugbee, 2013). Emission of low B from warm W LEDs can amplify stem elongation and leaf expansion, while high B from cold W LEDs can have the opposite effect, resulting in more compact specimens. Results from our preliminary experiment provide evidence that appropriate levels of R:Fr can greatly influence stem elongation to a greater degree than B (Magagnini et al., 2018). The reduction of photosynthetically active radiation resulting from shading limits the amount of R, B, and Fr received by the canopy, though the degree of R reduction tends to be far greater than that of Fr (Xu et al., 2020). Hence, low irradiance and wavelength perception work mutually to allow shoot elongation, perhaps in combination with the influence of exogenous sucrose. In agreement with these principles, the predicted optimal conditions for shoot elongation included low R:Fr ratios, including Fr intensities ranging from 30.6 – 43.3 μmol/m^2^/s and R between 0.3 – 19.1 μmol/m^2^/s. Likewise, shoot elongation was maximized under relatively low PAR light levels from 15.7 – 80.8 μmol/m^2^/s, while canopy area and number of nodes were predicted to be greater with low levels of Fr (1.8 – 24.7 μmol/m^2^/s and 8.8 – 19.9 μmol/m^2^/s, respectively) and higher PAR (139.1 – 194.5 μmol/m^2^/s, and 150.3 – 203.8 μmol/m^2^/s, respectively) fluence rates. As with previous literature, it appears that *in vitro* cannabis plants produce longer stems with fewer nodes and more narrow leaves when cultured at low light levels and higher amounts of Fr. These results demonstrate that *in vitro* cannabis plants respond to light signals similar to what would be expected *in vivo*. Further, the ability of machine learning and optimization algorithms to make predictions that agree with the general body of literature further supports the ability to recognize complex patterns using relatively quickly with few treatments. Thus, balances between alternative light spectra, their intensities and exogenously supplied carbohydrates are critical factors determining the outcome of many plantlet responses *in vitro*.

The effect of light quantity and quality studied together *in vitro* has been perused for many years in micropropagation but have been hampered due to the limitations of lighting systems and difficulties in proper replication (Kim et al., 2004; Miler et al., 2019; Tanaka et al., 1998). Many previous experiments explored the influences of single or binary combinations light spectra and their intensities on *in vitro* plantlet development (Lian et al., 2002; Manivannan et al., 2015; Shukla et al., 2017). Our study enlists novel LED technology combined with machine learning and optimization algorithms in an innovative system that assesses a vast assortment of sucrose concentrations and the cumulative impact of four different light qualities at a wide array of intensities to devise precision tissue culture protocols. Furthermore, for the first time, we suggest a superior machine learning and optimization algorithm approach for future plant tissue culture studies. Additionally, the results of the preliminary experiment exemplify that specific growth responses of *in vitro* cannabis can be directed by manipulating abiotic factors such as light intensity and quality in addition to exogenous carbon availability. This is further demonstrated by the results of the validation experiment. Such discoveries have valuable implications for the development of cannabis tissue culture techniques in the absence of PGRs.

## Conclusion

This machine learning –assisted, multivariable micropropagation study has demonstrated that distinct growth responses in cannabis can be shaped by changing the influences of sugar and light dynamics in the absence of PGRs. The development of alternative protocols to guide plant growth toward specific responses shows endless value for numerous *in vitro* applications. For instance, protocols to induce long stems, large internodes, many nodes, or many stems could be implemented when growing cultures for clonal propagation and sub-culturing, while cultures developing large root masses and large canopies could very well be more suited for *ex vitro* transfer. In addition, culmination of the protocols devised could be implemented, perhaps to trigger different developmental responses during different growth phases. Finally, the results obtained from this experiment allows us to recommend GRNN-SOS to be a more efficacious algorithm to study dynamic plant responses to multivariable stimuli *in vitro* for development of new methods, and optimization of current protocols. Rather than using traditional statistics to evaluate large datasets for making optimization predictions for tissue culture applications, the use of effective machine learning strategies for optimization of *in vitro* protocols should further be assessed as an alternative, or in combination with traditional statistical approaches to allow precision tissue culture practices.

## Conflict of Interest

The authors declare that the research was conducted in the absence of any commercial or financial relationships that could be construed as a potential conflict of interest.

## Author Contributions

MP, MH, and AMPJ conceived and designed the study, MP and MH conducted the experiments and analysed the data, AMPJ administrated the project and acquired funding, MP, MH, FS, and AMPJ wrote the manuscript. All authors approved the manuscript for publication.

## Funding

Funding for this project was provided through an NSERC discovery grant (RGPIN-2016-06252) awarded to AMPJ.

## Data Availability Statement

All processed data are available within the manuscript.

## Acknowledgments

Not applicable.

## Notes

### Competing Interest Statement

The authors have declared no competing interest.

